# Neuronal avalanches in awake macaque V1 and V4

**DOI:** 10.64898/2026.10.11.758173

**Authors:** Ksenia Solovyeva, Pavel Popov, Anna Makarova, Mikhail Lebedev

## Abstract

Neuronal avalanches are cascades of synchronised activity whose sizes follow power law. Analyses of avalanches shed light on cortical dynamics operating near a critical point. In awake monkeys, avalanches were found in motor and premotor cortex, where they are invariant to temporal binning, detection threshold and the number of electrodes employed. Whether awake primate early visual cortex, which receives strong structured afferent drive, shares these properties has not been tested. We analyzed a public 1,024-channel resting-state dataset from V1 and V4 of two macaques. In both animals avalanche sizes followed power laws over more than 10-fold range, the exponent scaled with bin width as α ∝ Δ*t*^β^ (β = −0.045 and −0.031), and the distributions ended at a cutoff proportional to the number of electrodes analysed. Raising the detection threshold from 1.5 to 4.5 standard deviations (SDs) removed over 99% of events but changed α by only 0.02–0.05 per SD, whereas randomly removing the same events destroyed the power law. Per-electrode shuffling abolished all heavy tails. Eye closure shifted α, β and the branching ratio by less than predefined equivalence bounds, far below the reproducible differences between recording sites. Avalanche organisation therefore extends to awake early visual cortex.

## Introduction

Cortical networks display spontaneous activity organised as oscillations, synchrony and travelling waves. Neuronal avalanches constitute a distinct motif in which activity propagates through cascades of varying sizes without a characteristic scale. Beggs and Plenz first showed in cortical slices and cultures that cascades of negative local field potential (LFP) deflections follow a power law in size with an exponent near −3/2, and that the branching ratio σ, the mean number of descendant activations per ancestor, lies close to unity: the signature of a critical branching process [1]. Models and experiments link this regime to maximal dynamic range and information transmission: for σ < 1 cascades die out, for σ > 1 they saturate the network, and only near σ ≈ 1 do propagation and stability balance [2–4].

In awake primates, Petermann et al. provided the benchmark demonstration in primary motor and dorsal premotor cortex [5]. Their argument rests on invariances rather than on a single power-law fit, because heavy-tailed histograms can arise from mechanisms unrelated to criticality [6]. The size distribution retains its form across temporal bin widths of 2–32 ms, with the exponent scaling as α ∝ Δ*t*^β^; the cutoff of the distribution tracks the number of electrodes analysed, as expected when the largest cascade is limited by the observed system rather than by the dynamics; and the organisation is unchanged when the detection threshold is raised from 1.5 to 4.5 SD, whereas randomly deleting the same number of events destroys it. Time-shuffled surrogates abolish the heavy tails.

Neuronal avalanches have subsequently been observed in the visual cortex of anaesthetized cats and in ex vivo turtle preparations [7,8]. However, it remains unknown whether the early visual cortex of awake primates exhibits the same dynamical regime and associated scaling invariances. This question needs careful investigation.

V1 receives strong, retinotopically organized feedforward input, whereas V4 occupies a higher level of the visual hierarchy and is characterized by larger receptive fields and distinct intrinsic timescales. If avalanche statistics are modulated by afferent input, these differences may be reflected in the dynamics of the two areas, particularly when retinal input is reduced during eye closure.

The dataset of Chen et al. [9] makes the test possible: 1,024 chronically implanted electrodes per animal spanning V1 and V4, resting-state sessions of 21–42 min with pupil-based labelling of eyes-open and eyes-closed epochs, per-channel signal-to-noise measurements from a same-day visually evoked task, and a documented cross-talk removal procedure. The 64-electrode arrays match the electrode counts of the original study and permit its spatial-subsampling test.

We asked (i) whether spontaneous LFP activity in V1 and V4 of awake macaques forms avalanches with the statistics and the three invariances reported for motor cortex, and (ii) whether these statistics depend on eye state, cortical area and recording site.

## Results

### Data selection and channel yield

Three resting-state sessions per animal were analysed (Table 1). Channels with a signal-to-noise ratio (SNR) below 2 in the same-day checkerboard session were discarded, as recommended by the data authors [9]; this kept 977–992 of 1,024 electrodes in monkey L and 361–411 in monkey A, in line with the counts reported for these sessions. Hardware cross-talk, present in all resting sessions of monkey L and essentially absent in monkey A, was handled by removing the first 50 entries of the data authors’ ranked removal list, the electrodes with the strongest synchrofact participation (Methods; the consequences of this choice are quantified below). Avalanches were analysed separately in every 8 × 8 array that retained at least 32 electrodes. In monkey L all 16 arrays qualified in every session (37–64 electrodes, median 60). In monkey A, whose implant had lower signal quality at the time of recording, seven V1 arrays qualified (19 array–sessions; 34–64 electrodes, median 53); both V4 arrays of this animal retained at most 12 electrodes and were excluded, so comparisons between V1 and V4 rest on monkey L (Supplementary Fig. S1).

**Table 1.** Sessions, electrode yield and condition durations. ^a^Electrodes with SNR ≥ 2 across the 1,024-electrode implant in the same-day checkerboard session, as reported by the data authors [9]; our join of the same files retained 361, 411 and 378 in monkey A. ^b^Arrays with ≥ 32 retained electrodes, and the median number of retained electrodes per analysed array. ^c^After slow-wave exclusion; medians across arrays.

| Animal | Session | Date | Duration | SNR $\geq 2^a$ | Arrays<br>(electrodes)<sup>b</sup> | Eyes open<br>(s)<sup>c</sup> | Eyes<br>closed (s)<sup>c</sup> |
| --- | --- | --- | --- | --- | --- | --- | --- |
| L | L_RS_250717 | 25/07/2017 | 22 min 42 s | 981 | 16 (60.5) | 354 | 824 |
| L | L_RS_090817 | 09/08/2017 | 22 min 00 s | 977 | 16 (58.5) | 301 | 837 |
| L | L_RS_100817 | 10/08/2017 | 21 min 37 s | 992 | 16 (59.0) | 266 | 821 |
| A | A_RS_140819 | 14/08/2019 | 32 min 34 s | 359 | 5 (63) | 1,844 | 48 |
| A | A_RS_150819 | 15/08/2019 | 38 min 17 s | 416 | 7 (53) | 2,100 | 157 |
| A | A_RS_160819 | 16/08/2019 | 42 min 00 s | 379 | 7 (52) | 2,156 | 230 |

Periods dominated by slow waves were removed with the spectral criterion of the original study [5] (Methods), which excluded 8.5–13.0% of eyes-open and 14.6–17.3% of eyes-closed time in monkey L. Eye state was strongly imbalanced, in opposite directions in the two animals (Table 1): monkey L spent 70–77% of each session with eyes closed, monkey A 90–97% with eyes open, consistent with the proportions reported by the data authors for the same sessions. The eye-state contrast was therefore restricted to monkey L (266–354 s eyes open and 821–837 s eyes closed per session after exclusion); monkey A, with 48–230 s of eyes closed per session, contributes whole recordings.

### Avalanche sizes follow power laws in both animals

Negative LFP deflections (nLFPs) occurred in spatiotemporally clustered patterns rather than independently across electrodes (Fig. 1). Pooled over arrays and sessions, monkey L yielded 38,661 (eyes open) and 71,351 (eyes closed) avalanches at Δ*t* = 2 ms and 7,955 and 17,050 at 32 ms; monkey A yielded 68,302 and 20,389 (Table 2). In every condition and at every bin width, sizes followed a power law over more than a 10-fold range (decade) (Fig. 2a), and the power law was favoured over an exponential (normalised log-likelihood ratio *R* = 12–86 in monkey L and 66–110 in monkey A). Maximum-likelihood exponents ranged from α = −1.71 at Δ*t* = 2 ms to −1.49 at 32 ms in monkey L, eyes-open and eyes-closed values lying within 0.02 of each other at every bin width, and from −1.82 to −1.67 in monkey A (Table 2). Bootstrap resampling of avalanches gave 95% confidence intervals of ±0.004–0.009 for these pooled exponents; the uncertainty of a group mean is dominated instead by differences between arrays (Supplementary Table S1).

**Figure 1.**
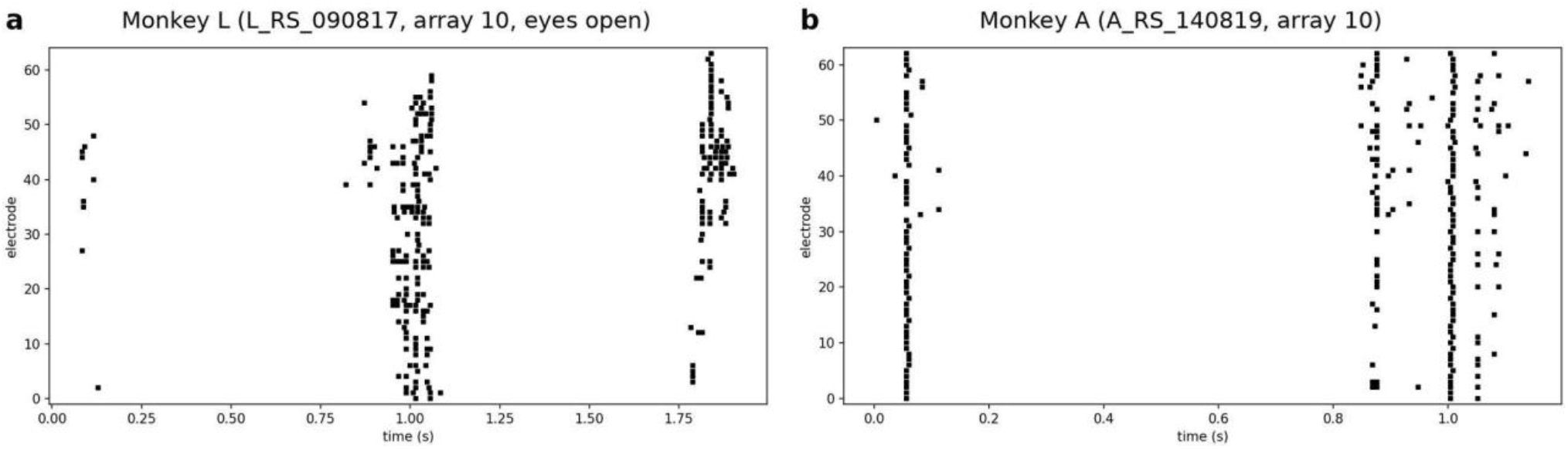
Spontaneous nLFPs cluster into avalanches. nLFP rasters at Δ*t* = 4 ms in a 2-s window centred on the largest avalanche of the recording. (**a**) Monkey L, session L_RS_090817, array 10, eyes open (largest avalanche 97 nLFPs). (**b**) Monkey A, session A_RS_140819, array 10 (largest avalanche 62 nLFPs). Each dot is one supra-threshold negative LFP deflection on one electrode; avalanches are runs of consecutive non-empty bins.

**Figure 2.**
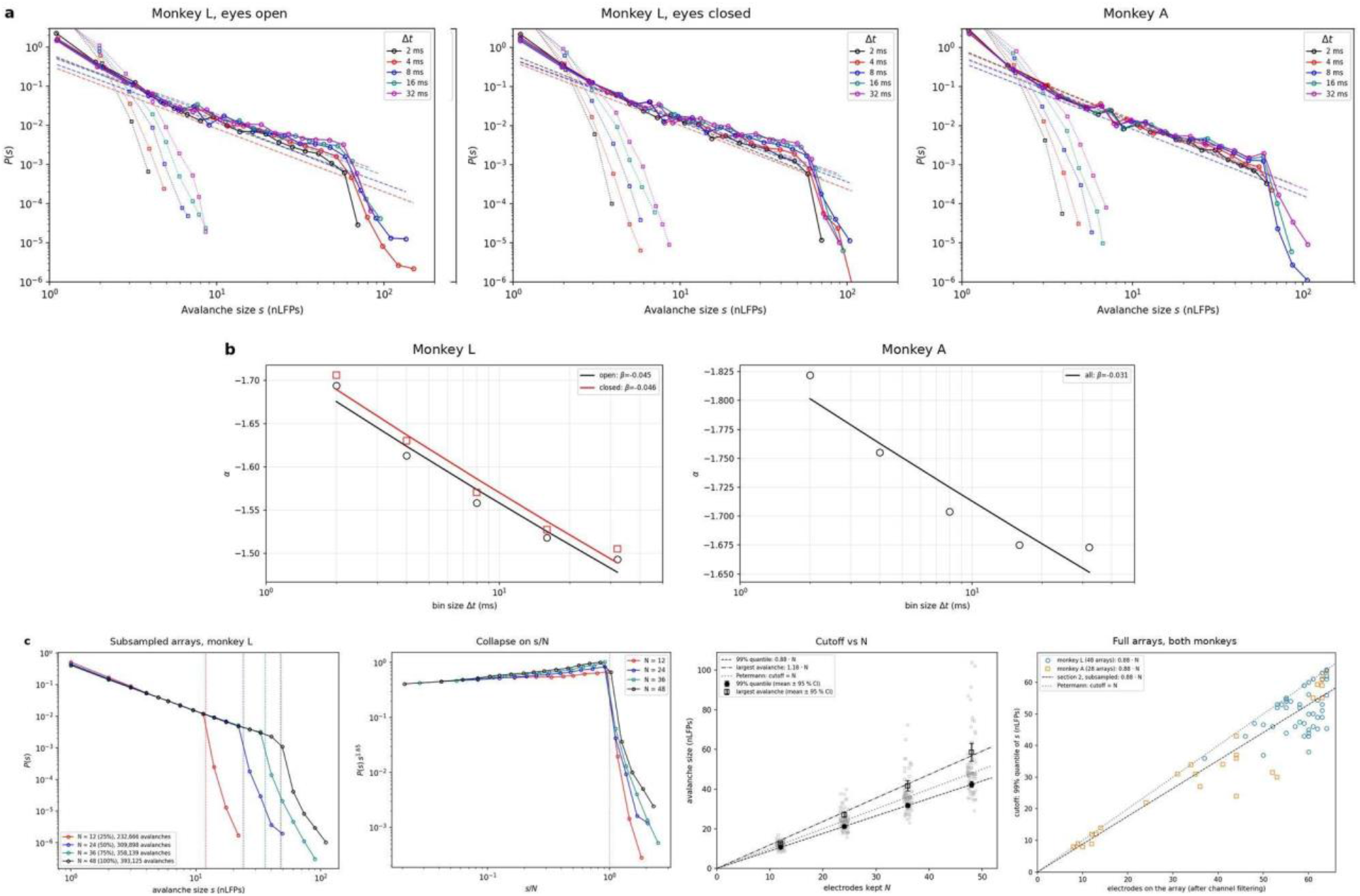
Size distributions, their dependence on bin width and on the number of electrodes. (**a**) Avalanche size distributions for five bin widths (2 ms black, 4 ms red, 8 ms blue, 16 ms teal, 32 ms magenta), pooled across arrays and sessions: monkey L eyes open and eyes closed, monkey A whole recordings. Circles and solid lines: data; dashed lines: maximum-likelihood power-law fits (*x*_min_ = 1); squares and dotted lines: per-electrode time-shuffled surrogates. (**b**) Size exponent α of the pooled distributions against bin width, ordinate inverted as in the original study; lines are fits of α ∝ Δ*t*^β^. (**c**) Finite-size scaling (Δ*t* = 4 ms, whole recordings). From left: *P*(*s*) for random subsets of *N* = 12, 24, 36 and 48 electrodes in monkey L (46 array–sessions, five subsets each; dotted lines at *s* = *N*); the same distributions as *P*(*s*)·*s*^1.65^ against *s*/*N*; cutoff (99th percentile, filled) and largest avalanche (open) against *N*, grey points single array–sessions, black mean ± 95% CI, lines proportional fits, dotted line cutoff = *N*; cutoff against the retained electrode count of full arrays in both animals.

Shuffling each electrode’s binned events in time, which preserves its rate but destroys the timing across electrodes, converted every distribution into a rapidly decaying one (Fig. 2a, squares). In monkey L, the shuffled distributions had apparent exponents of −2.7 to −5.2 and were better described by an exponential at every bin width (*R* = −27 to −57).

### Temporal scale invariance

The magnitude of α decreased monotonically with bin width (Fig. 2b), and log|α| was linear in log Δ*t* over 2–32 ms, giving β = −0.045 (eyes open) and −0.046 (eyes closed) for the pooled distributions of monkey L and −0.031 for monkey A. Fitted per array–session, β in monkey L had a median of −0.045 (interquartile range −0.050 to −0.036 eyes open, −0.049 to −0.038 eyes closed); none of the 96 estimates was positive, and 67% fell inside the range reported for M1 and PMd (−0.16 to −0.04) [5]. In monkey A, α flattened between 16 and 32 ms (−1.675 and −1.673), which placed β just outside that range (per-file median −0.030, range −0.045 to −0.011, 3 of 19 inside the M1/PMd range); in a mixed model the difference between the animals was 0.015 (95% CI 0.010–0.020; Supplementary Table S2).

β depended on which bin widths entered the fit (Supplementary Fig. S2): in monkey L its median was −0.067 for Δ*t* = 2–4 ms, −0.045 for 2–32 ms and −0.029 for 8–32 ms, with 96%, 67% and 18% of files inside the M1/PMd range. We report the full 2–32 ms range, as in the original study; values of β are comparable across studies only for matched sets of bin widths.

Because |β| is small, the distributions for different bin widths already overlapped over their power-law range without any rescaling (Fig. 2a). The rescaling used in the original study, *P*(*s*)·Δ*t*^0.22^ against *s*·(Δ*t*/4 ms)^−β^, did not collapse them when applied with our fitted β: its amplitude factor separated the curves by up to a factor of three in both animals (Supplementary Fig. S3). The amplitude exponent of 0.22, obtained for motor cortex, therefore did not transfer to these data.

### The cutoff scales with the number of electrodes

At criticality the largest avalanches are bounded by the size of the observed system rather than by the dynamics, so the cutoff of *P*(*s*) should move in proportion to the number of electrodes analysed [5]. The pooled distributions ended in a shoulder close to the number of retained electrodes per array (Fig. 2a). To test proportionality directly, we drew five random subsets of *N* = 12, 24, 36 and 48 electrodes from each of the 46 array–sessions of monkey L that retained at least 48 electrodes (Δ*t* = 4 ms, whole recordings) and rebuilt the avalanches. The cutoff, taken as the 99th percentile of sizes, was 10.7, 21.2, 31.7 and 42.4 nLFPs — a constant 0.88 *N* — and the largest avalanche stayed between 1.07 *N* and 1.22 *N* (Fig. 2c). Fitting cutoff ∝ *N*^D^ per array–session gave *D* = 0.99 ± 0.03 (mean ± SD), close to but statistically distinguishable from 1 (Wilcoxon *P* = 0.035). Rescaled by *N*, the subsampled distributions collapsed onto one curve.

The arrays’ natural differences in retained channel count provided an independent check without subsampling. Across full arrays the cutoff tracked the number of retained electrodes in both animals, most clearly in monkey A, whose arrays spanned 8–64 electrodes (*r* = 0.95, *P* = 8 × 10^−15^; monkey L, 37–64 electrodes, *r* = 0.53, *P* = 1 × 10^−4^), with cutoffs of 0.88 ± 0.13 and 0.89 ± 0.11 times the electrode count.

### Threshold invariance

The third invariance concerns the amplitude axis. We re-detected nLFPs at thresholds *z* from 1.5 to 4.5 SD in steps of 0.5 SD, keeping electrodes and per-electrode statistics fixed (Δ*t* = 4 ms, whole recordings, 67 array–sessions). Raising the threshold from 1.5 to 4.5 SD reduced the nLFP rate 164-fold in V1 and 143-fold in V4 of monkey L and 127-fold in V1 of monkey A (Fig. 3a). The size distributions nonetheless kept their power-law form at every threshold (Fig. 3b; *R* > 0 throughout; Supplementary Fig. S4, Supplementary Table S3) and α changed little (Fig. 3c): from −1.54 to −1.65 in V1 and from −1.52 to −1.59 in V4 of monkey L, and from −1.63 to −1.83 in V1 of monkey A. Per array, the slope dα/d*z* was 0.033 ± 0.045 per SD in V1 and 0.024 ± 0.033 in V4 of monkey L and 0.049 ± 0.116 in monkey A (mean ± SD), within the 0.07 ± 0.06 per SD reported for motor cortex [5]. This small drift was detectable across the 42 V1 array–sessions of monkey L (Wilcoxon test against zero, *P* = 9 × 10^−6^) but not in the smaller groups (V4, *P* = 0.44; monkey A, *P* = 0.14). Mean avalanche size changed by at most a quarter (from 9.5 to 7.2 nLFPs in V1 and 11.4 to 11.3 in V4 of monkey L; 7.4 to 6.7 in monkey A).

**Figure 3.**
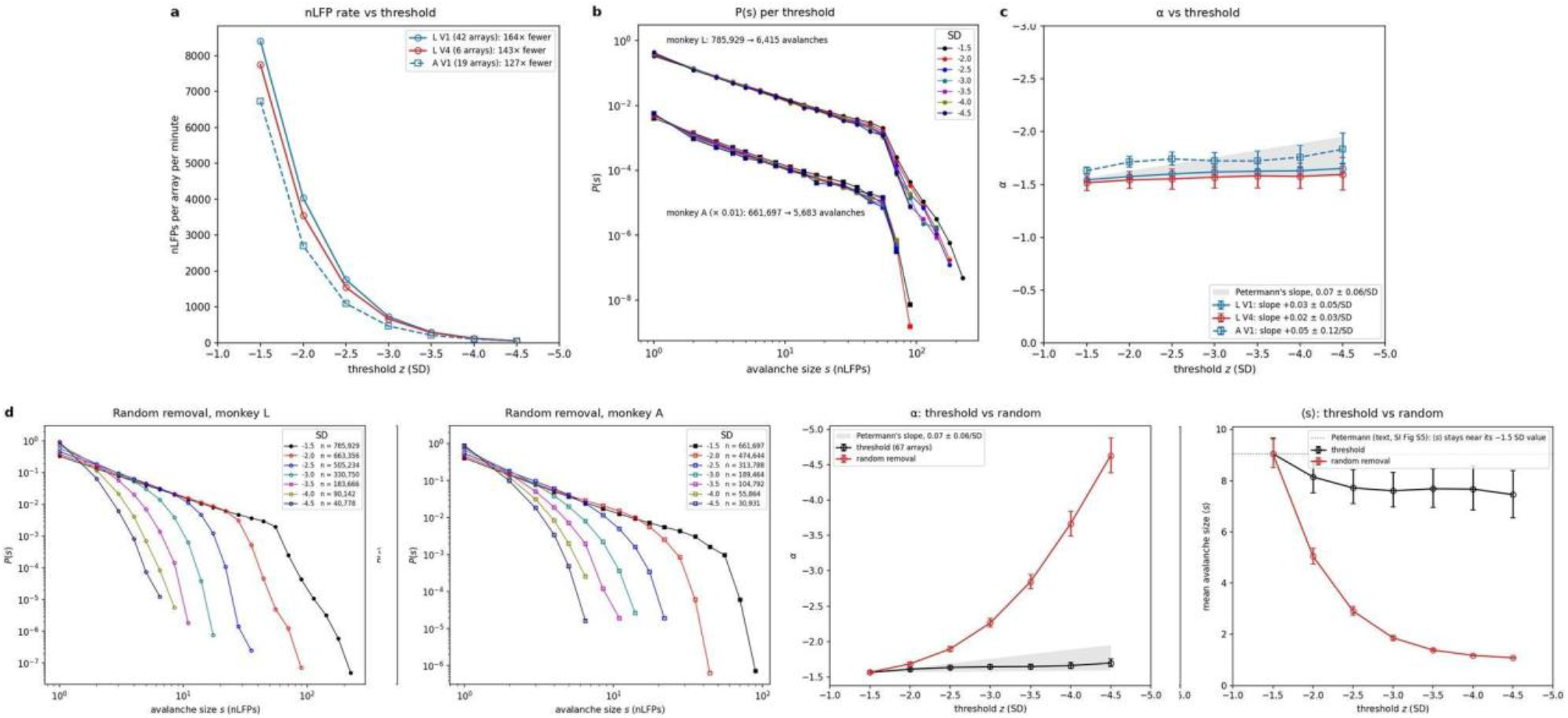
Threshold invariance and the random-removal control. Δ*t* = 4 ms, whole recordings; one line per animal × area, means over array–sessions (monkey L: 42 V1, 6 V4; monkey A: 19 V1). (**a**) nLFPs per array per minute against threshold *z*. (**b**) *P*(*s*) at each threshold from −1.5 to −4.5 SD, pooled per animal (monkey A shifted down by a factor of 100 for clarity). (**c**) α against *z* with 95% bootstrap confidence intervals over arrays; the grey band marks the slope range reported for motor cortex. (**d**) Random removal: *P*(*s*) at −1.5 SD (filled) and after deleting events at random down to the count of each higher threshold (open), and α and mean size ⟨*s*⟩ for threshold-based versus random removal at matched event counts.

Threshold-based thinning was not equivalent to random thinning (Fig. 3d). When the 1.5-SD events of each array were deleted at random down to the count obtained at each higher threshold, the distributions steepened sharply (α = −4.7, −4.3 and −4.5 at the 4.5-SD count in the three groups), mean size fell towards one nLFP (1.07–1.11), and an exponential was favoured from the 2.5-SD count onward. The random-removal slope dα/d*z* was about 1 per SD in every group (1.04 ± 0.32, 0.91 ± 0.22 and 0.98 ± 0.46), 20–40 times the threshold slope (paired Wilcoxon test over 67 array–sessions, *P* = 1 × 10^−12^). Large avalanches are therefore not random aggregates of small ones: the events that survive a high threshold remain organised into the same scale-free cascades.

### Avalanche lifetimes

Lifetime distributions were heavy-tailed at all bin widths and were abolished by shuffling (Fig. 4b). Expressed in multiples of Δ*t*, the distributions for 2–8 ms overlapped, whereas those for 16 and 32 ms ended earlier. At Δ*t* = 2 ms the lifetime exponent was τ = −2.14 (eyes open) and −2.17 (eyes closed) in monkey L and −2.32 in monkey A, steepening with bin width to −2.75, −3.03 and −3.37 at 32 ms, where most avalanches span only a few bins (Table 2). Size and lifetime were only loosely related (Fig. 4a): large avalanches occurred at every lifetime, and at Δ*t* = 4 ms some avalanches recruited 40 or more nLFPs within one or two bins — near-synchronous events across most of the array (Fig. 1b).

**Figure 4.**
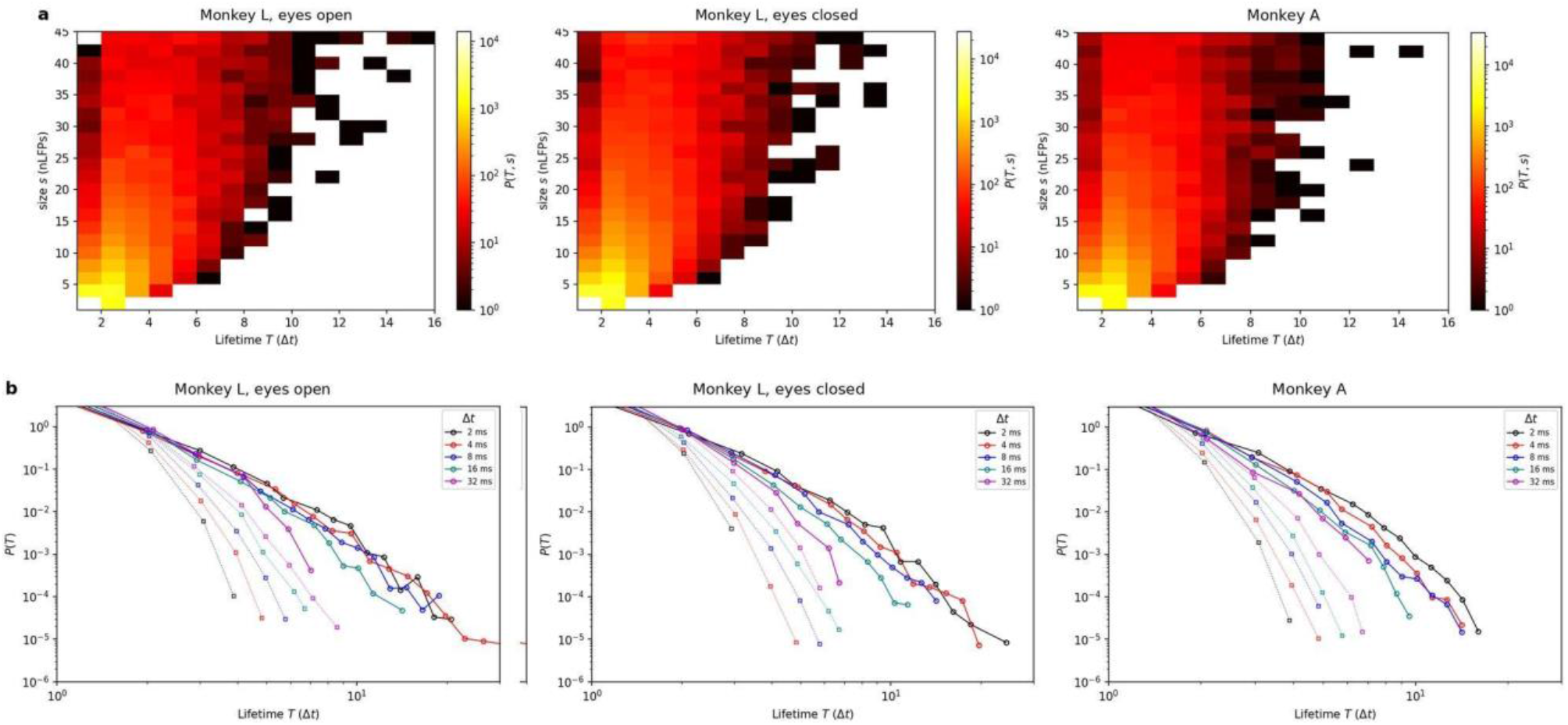
Avalanche lifetimes. (**a**) Joint distribution of avalanche size and lifetime at Δ*t* = 4 ms (monkey L eyes open and eyes closed; monkey A whole recordings); colour, number of avalanches (logarithmic scale). (**b**) Lifetime distributions, with lifetime in multiples of Δ*t*, for five bin widths; circles and solid lines, data; squares and dotted lines, per-electrode shuffled surrogates. Colour code as in fig. 2.

### Branching ratio

We computed three estimators of σ on identical binned rasters (Methods; Fig. 5, Supplementary Table S4). They differed in how the end of an avalanche is counted, and that difference dominated their values. The within-avalanche estimator, which ignores terminations, stayed at 0.99–1.02 at every bin width. The estimator that counts every transition, including the one into the empty bin that ends an avalanche, fell from 0.67 at 2 ms to 0.25–0.32 at 32 ms: wider bins compress avalanches into fewer bins, so the terminating zero carries proportionally more weight. The Beggs–Plenz estimator [1,5], which counts the first transition of each avalanche, gave σ = 0.95 ± 0.28 (eyes open) and 0.96 ± 0.28 (eyes closed) at Δ*t* = 2 ms (mean ± SD over 48 array–sessions) and decreased with bin width in the same way. Per-electrode shuffling reduced the terminating-transition estimator from 0.67 to 0.02–0.03 at 2 ms (0.14–0.18 at 32 ms), so the near-unity values do not follow from the event rate alone. Monkey A showed the same pattern (Fig. 5): the Beggs–Plenz estimator was 1.04 ± 0.38 at Δ*t* = 2 ms, the within-avalanche estimator 0.90–0.99 across bin widths, and the terminating-transition estimator fell from 0.64 to 0.22, 0.044 below monkey L at Δ*t* = 4 ms (95% CI 0.012–0.075).

**Figure 5.**
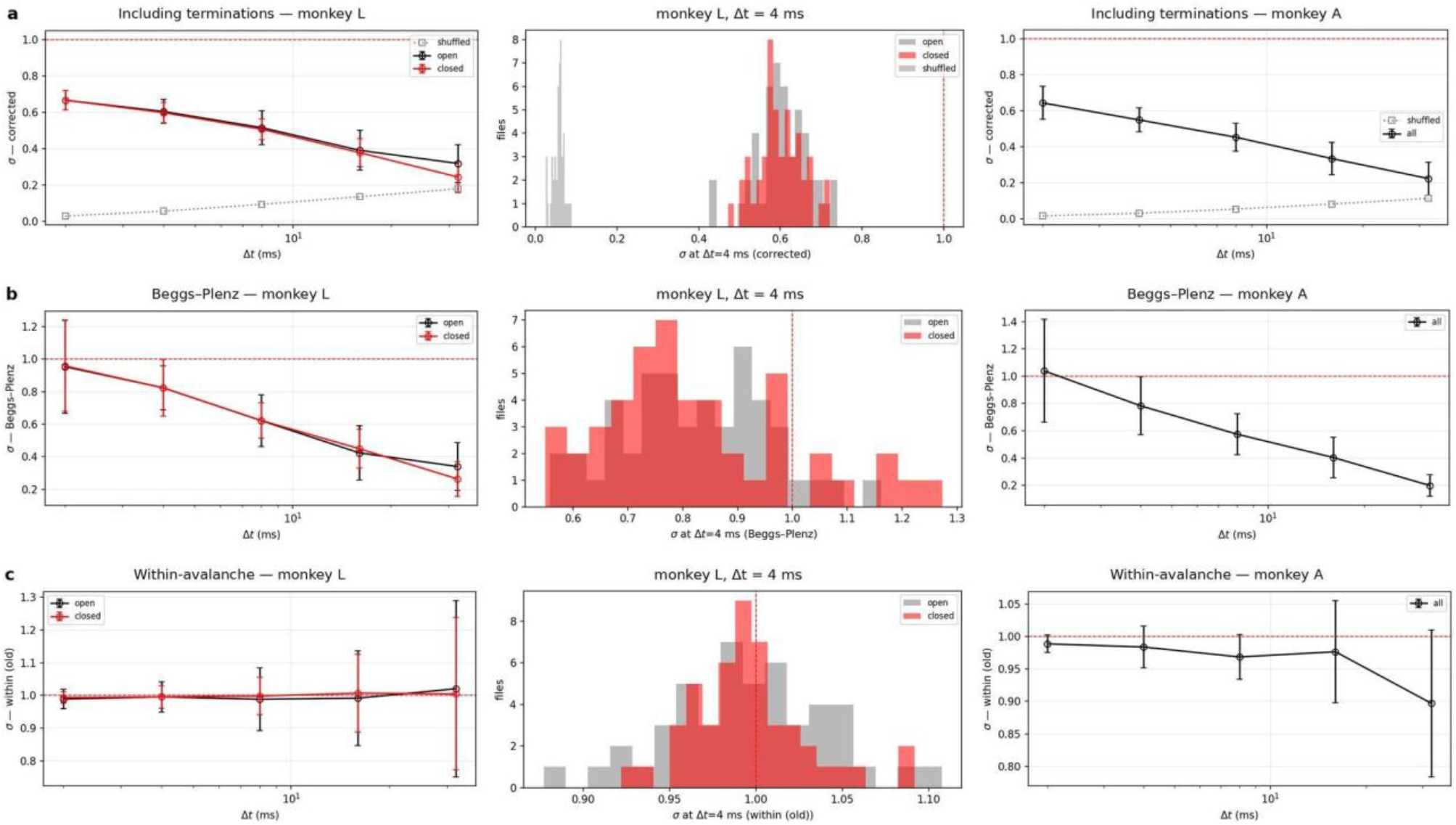
The branching ratio depends on the estimator and the bin width. Rows: (**a**) the estimator including terminations, (**b**) the Beggs–Plenz estimator, (**c**) the within-avalanche estimator. Left, σ against Δ*t* in monkey L (mean ± SD over 48 array–sessions per eye state; grey squares in **a**, per-electrode shuffled surrogates); middle, distribution across array–sessions of monkey L at Δ*t* = 4 ms; right, σ against Δ*t* in monkey A (mean ± SD over 19 array–sessions). Dashed lines: σ = 1.

### Eye state leaves the avalanche parameters unchanged

In monkey L the same electrodes were recorded in both eye states in 48 array–sessions, which allowed a paired comparison (Fig. 6a). Mean α at Δ*t* = 4 ms was −1.607 with eyes open and −1.616 with eyes closed, σ (terminating-transition estimator) 0.605 and 0.599, and β −0.043 and −0.045. In a linear mixed model with arrays nested in sessions, none of the main effects of eye state was significant (eyes closed minus open: α −0.005, 95% CI −0.013 to 0.003, *P* = 0.23; σ −0.005, −0.018 to 0.009, *P* = 0.47; β −0.001, −0.004 to 0.002, *P* = 0.40). Paired tests at each bin width, corrected for multiple comparisons, were likewise null with one exception: at Δ*t* = 32 ms σ was lower with eyes closed (−0.074, *P*_BH_ = 3 × 10^−7^; Supplementary Table S5), which appears in the model as an eye state × bin width interaction for σ (−0.016 per doubling of Δ*t, P* = 9 × 10^−5^) and was absent for α.

**Figure 6.**
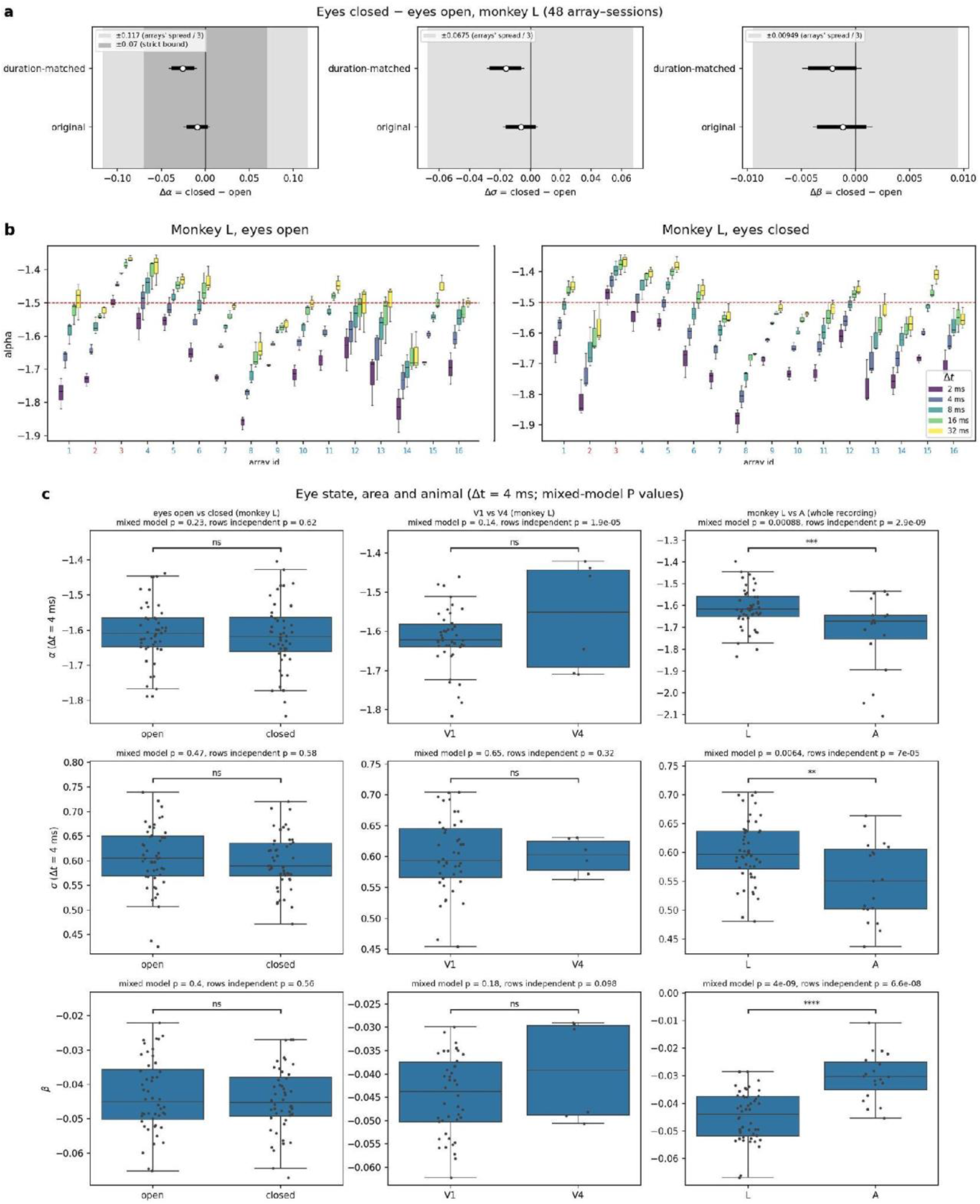
Eye state, recording site, area and animal. (**a**) Eyes closed minus eyes open in monkey L for α and σ at Δ*t* = 4 ms and for β: mean paired difference over 48 array–sessions with 90% and 95% bootstrap confidence intervals, for the original and the duration-matched comparison; shaded, equivalence bounds. (**b**) α per array in monkey L, eyes open and eyes closed, at five bin widths (colour); each box spans the three sessions; V4 array numbers in red; dashed line, α = −1.5. (**c**) α and σ at Δ*t* = 4 ms and β for eyes open against eyes closed (monkey L), V1 against V4 (monkey L) and monkey L against monkey A (whole recordings), one point per array–session; brackets give the mixed-model *P* value, panel titles also the *P* value obtained when rows are treated as independent.

To distinguish a lack of statistical significance from evidence of equivalence, we performed two one-sided tests [10] using bounds defined as one third of the range of each parameter across arrays (α = 0.117, σ = 0.068, β = 0.009), with a stricter bound of 0.07 applied for α. The paired differences lay inside every bound (all *P*_TOST_ < 10^−6^); for α the 95% CI of the difference was −0.024 to 0.005. Eyes-closed recordings were 2.3–3.1 times longer than eyes-open ones, so we recomputed the eyes-closed parameters on five random windows per array matched to the eyes-open duration of the same array. Matching revealed small systematic shifts in α and σ, with α increasing by 0.026 when the eyes were closed (95% CI 0.010–0.041; Wilcoxon P = 0.004) and σ decreasing by 0.016 (0.004–0.028; P = 0.012), whereas β remained unchanged (P = 0.27). All observed shifts remained within the equivalence bounds (all *P*_TOST_ ≤ 2 × 10−^6^). Eyes-closed epochs that survived the slow-wave exclusion were still slower than eyes-open ones: in one example array per session, the median fraction of LFP power in 1–10 Hz was 0.82 in retained eyes-closed windows against 0.71–0.74 with eyes open (Supplementary Fig. S5).

### Avalanche statistics differ between recording sites more than between conditions

The largest difference in the parameter estimates lay between arrays. In monkey L, α at Δ*t* = 4 ms was reproducible for a given array across the three sessions but differed between arrays, with per-array medians from −1.44 to −1.81 (Fig. 6b). In the mixed model, arrays accounted for 83% of the variance of α in monkey L (between-array SD 0.082 against a within-array residual SD of 0.037) and 95% when both animals were combined; for σ and β the shares were 41% and 53%. The largest eye-state difference (0.026 in α after duration matching) is thus about a third of the between-array SD, which is why eye state was compared within arrays.

Comparing areas within monkey L (2 V4 against 14 V1 arrays, three sessions each), we found no detectable difference: V4 minus V1 was 0.053 for α (95% CI −0.018 to 0.124, *P* = 0.14), −0.011 for σ (*P* = 0.65) and 0.005 for β (*P* = 0.18; Fig. 6c). Treating every array–session–bin-width row as an independent observation would instead have given *P* = 2 × 10^−5^ for α, illustrating how pseudo-replication inflates contrasts that are defined between arrays. The two animals did differ: in monkey A α was steeper by 0.107 (95% CI 0.044–0.170, *P* = 9 × 10^−4^), σ lower by 0.044 (*P* = 0.006) and β shallower by 0.015 (*P* = 4 × 10^−9^). With three sessions per animal these differences describe the two individuals rather than a population; part of the difference in α may be technical, since monkey A’s arrays retained fewer electrodes (median 53 against 60) and electrode removal alone steepens α (next section).

### Robustness to preprocessing choices

#### Cross-talk

We recomputed all parameters in monkey L with none of the cross-talk list, its first 50 entries (the setting used throughout) and the full list (186–250 electrodes per session), and, as a control, after removing as many randomly chosen electrodes as the full list (Supplementary Fig. S6). Mean α at Δ*t* = 4 ms was −1.605, −1.613, −1.658 and −1.643 respectively, σ 0.604, 0.601, 0.574 and 0.564, and β −0.044 to −0.048. Removing the full list (on average 11 more electrodes per array than the default) steepened α by 0.045, but removing the same number of random electrodes steepened it by 0.030 and lowered σ even more than the list did. The part of the change attributable to cross-talk itself is therefore about 0.015 in α and none in σ; the remainder is the finite-size effect of observing fewer electrodes.

#### Detection and binning

Results were recomputed for four combinations of event definition (one event per threshold excursion, or every local minimum below threshold) and binning (detection on the decimated signal, or detection at 2 ms followed by merging into wider bins; Methods). In monkey L, α at Δ*t* = 4 ms varied by less than 0.04 across combinations, and the median β over 2–32 ms ranged from −0.032 to −0.054; no file in any combination yielded a positive β (Supplementary Fig. S7). Averaging the size distributions over all *n* possible start phases of the decimation, as in the original study, changed α by at most 0.021 (mean ≤ 0.002) and β by 0.0004 ± 0.0022 on the 16 arrays of one session (Supplementary Fig. S8), so a single phase was used throughout. Counting every below-threshold sample as an event, instead of one event per excursion, would have inflated the event count 9.9-fold (interquartile range 8.6–11.5) at 3 SD (Supplementary Fig. S9).

## Discussion

Applied to V1 and V4 data obtained in awake macaques, a pipeline built to the specification of the original study [5] returned similar results in both animals: power-law size distributions spanning more than a decade with exponents between −1.8 and −1.5 depending on bin width, heavy-tailed lifetimes, scaling of the exponent with bin width, a cutoff proportional to the number of electrodes observed, and invariance of the size distribution to a more than hundredfold change in the number of detected events. Surrogates that preserve single-electrode rates but destroy cross-electrode timing abolished the heavy tails, and random thinning at matched event counts destroyed the power law that threshold-based thinning preserves. On these measures, sensory cortex under strong structured afferent drive did not exhibit patterns distinguishable from that of premotor and motor cortex, which supports power-law avalanches as a general property of cortical networks.

Because power-law-shaped histograms could arise from many mechanisms [6], the weight of the argument rests on the invariances. Three invariances were present here, each with a quantitative departure that should be stated. Temporal invariance held, but β depended on the set of bin widths fitted, the amplitude rescaling of the original collapse did not carry over, and, in monkey A, the flattening of α between 16 and 32 ms placed β (−0.031) slightly outside the range reported for motor cortex. The dependence of β on the bin widths is one instance of a general problem: statistics inferred from neural time series depend on the rate at which the system is sampled, coarser sampling can change rather than merely blur the structure recovered from the data, and analysing several sampling rates together can be exploited rather than only corrected for [16,17]. Spatial invariance held in the strong form of a cutoff proportional to electrode count (*D* = 0.99) in subsampled arrays and in the natural variation of channel count across arrays; random rather than contiguous subsampling, which we used to avoid distortion of avalanche distributions [11]. The near-linear scaling we observed was reassuring.

Threshold invariance held in the sense of the original study (a slope of α on threshold of a few hundredths per SD against about 1 per SD for random thinning) but with 42 V1 array–sessions the residual drift of 0.03 per SD became statistically detectable. We consider this as a mild deviation from exact amplitude invariance that the original study, with fewer arrays, did not resolve.

On identical rasters our three estimators spanned 0.25 to 1.02, and the spread followed from whether the terminating transition of an avalanche was counted and from how the bin width compared with the interval between events. Reported values of σ are therefore not comparable across studies unless the estimator and the bin width are specified, and a value below unity should not by itself be read as evidence of subcritical dynamics. In addition, every estimator computed from avalanches was biased when only a small fraction of the network is observed [11,12]; estimators that are invariant to subsampling, such as multistep regression [12], suggested that cortex in vivo operates in a slightly subcritical, reverberating regime [13]. We therefore frame our result as branching consistent with unity under the estimator used in the source literature, not as a measurement of exact criticality.

Withdrawal of visual input during eye closure changed α, β and σ by less than predefined equivalence bounds across 48 matched array–session pairs. When durations were matched, a small shift emerged (α steeper by 0.026 and σ lower by 0.016 with eyes closed) in the direction of slightly more subcritical dynamics, but even this shift is a third of the between-array spread. Two interpretations are possible. The network could be held near its operating point across changes in afferent drive of this size, so that eye closure is too weak a perturbation to displace it. Alternatively, eye closure in this dataset could not isolate the intended variable: prolonged closure co-occurs with drowsiness, the eyes-closed periods that survive slow-wave exclusion are still slower than eyes-open ones, and the matched shift could equally reflect vigilance. Separating these requires an independent vigilance measure and, ultimately, a designed perturbation rather than a spontaneous one.

The clearest structure in our estimates was between recording sites rather than between conditions: α was reproducible for a given array across sessions and spans −1.44 to −1.81 across arrays of one animal, with arrays carrying 83–95% of its variance. Part of this is technical, including retained electrode count, residual artefact, position relative to the sulcus. The dependence of α on electrode count shown by the cross-talk control means that electrode count should be controlled before any site difference is interpreted. Neither V1 versus V4, tested on two V4 arrays in one animal, nor the eye-state contrast accounts for the spread. Relating per-array statistics to receptive-field position, which is available for this dataset [9], is a direct way to ask whether what remains reflects the functional architecture.

Utah arrays provide no laminar resolution, whereas the avalanche organisation described in the literature is specific to superficial layers; our estimates average over depth. Two animals with three sessions each, an eye-state contrast in one animal only and a V1–V4 comparison resting on two V4 arrays limit generalisation, and the resting condition carries no behavioural markers beyond eye state. Monkey L’s recordings are those most affected by hardware cross-talk, and electrode removal does not eliminate sporadic low-rate artefacts. We did not validate nLFPs against local spiking, although the dataset provides multi-unit envelope signals that would allow it, used a single surrogate type that preserves event rates rather than the LFP spectrum, did not test the spatial contiguity of the electrodes that take part in an avalanche against the prediction of volume conduction. This is relevant because some large avalanches recruit most of the array within one or two bins (Figs. 1b, 4a). We did not test the scaling relation between size and lifetime exponents or avalanche shape collapse [14,15], which postdate the original study.

The avalanche organisation previously documented in motor and premotor cortex thus extends to early visual cortex. V1 and V4 of two macaques produced power-law size distributions, heavy-tailed lifetimes, exponent scaling with bin width, cutoffs proportional to electrode count, and invariance to threshold, while shuffled surrogates and random thinning destroyed these signatures. We found quantitative departures: a shallower β in one animal, a statistically detectable but small drift of α with threshold, and the failure of the original amplitude rescaling. This suggests that the invariances are approximate rather than exact, and that their parameters are shaped in part by technical factors such as electrode count and array position. The absence of a detectable effect of eye closure on α, β, and σ, within equivalence bounds and against a between-array variance that dominates every condition contrast, suggests either that the dynamics are robust to withdrawal of retinal drive or that spontaneous eye closure is too weak, and too confounded with vigilance, to serve as a decisive perturbation. Whether the reproducible per-array differences we observed reflected the functional architecture of the recorded tissue or residual technical variability remains an open question, and answering it will require controlled electrode-count matching, validation of nLFPs against spiking and spatial contiguity, and, ultimately, designed perturbations that displace the network from its operating point and track its recovery.

## Methods

### Dataset

We analyzed the openly available resting-state recordings of Chen et al. [9] (G-Node Infrastructure, https://doi.org/10.12751/g-node.i20kyh). Two adult male rhesus macaques (*Macaca mulatta*, monkeys A and L), both 7 years old and weighing 11.0 and 12.6 kg at the time of cortical implantation, were chronically implanted with sixteen 8 × 8 Utah arrays (64 iridium-oxide electrodes each, 1.5 mm shank length, 400 µm inter-shank spacing) connected to a 1,024-channel pedestal. Fourteen arrays were placed in V1 and two in V4 of the left hemisphere, with receptive fields covering the central 6–9 degrees of visual angle in one quadrant of the visual field. Array-to-area assignment was taken from the channel–area maps supplied with the dataset (V4: arrays 2 and 3 in monkey L, arrays 2 and 5 in monkey A). At the time of the resting-state recordings the pedestal had been implanted for 3 months (monkey L) and 1 year (monkey A).

For the resting-state sessions the animal was seated and head-fixed in a darkened, quiet room, performed no task, and was free to shift gaze or fall asleep. Three sessions per animal were used (Table 1). All animal procedures were performed by the original authors in accordance with the NIH Guide for the Care and Use of Laboratory Animals and approved by the institutional animal care and use committee of the Royal Netherlands Academy of Arts and Sciences (approval AVD8010020171046) [9]. No new animal experiments were performed for the present study.

### Signals and preprocessing

Broadband signals were acquired at 30 kHz. We used the LFP supplied with the dataset (low-pass 150 Hz, fourth-order Butterworth, downsampled to 500 Hz, 50 Hz band-stop) and band-passed it at 1–100 Hz with a fourth-order Butterworth filter applied forward and backward (zero phase), matching the LFP band of the original study [5].

Channel quality was taken from the same-day checkerboard session of each resting-state session, in which the SNR is the peak stimulus-evoked multi-unit response minus mean spontaneous activity, divided by the standard deviation of spontaneous activity [9]; electrodes with SNR < 2 were discarded. Hardware cross-talk was addressed with the data authors’ ranked removal list, derived from near-simultaneous threshold crossings across electrodes (synchrofacts) tested against dithered surrogates [9,18]. Electrodes enter the list in decreasing order of synchrofact participation; we removed the first 50 entries of each session’s list (about three electrodes per array), over which participation falls from 0.35–0.85 to 0.07–0.23, whereas the remainder of the list (up to 250 electrodes) continues down to below 0.01.The effect of this choice is quantified in Results. Arrays with fewer than 8 retained electrodes were not processed, and all analyses reported here use arrays with at least 32.

Periods of sleep or drowsiness were removed with the criterion of the original study [5]: in 2-s windows advanced in 1-s steps, the fraction of each electrode’s power in 1–10 Hz was computed (Welch’s method), and a window was discarded when more than 50% of the array’s electrodes had more than 90% of their power in that band. Eye state was taken from the pupil-based segmentation supplied with the dataset, in which pupil diameter is baseline-corrected, downsampled to 1 Hz and thresholded [9]. For each condition (the full awake recording, eyes open, and eyes closed), the retained samples were concatenated, and conditions with less than 10 s of data were excluded from the analysis.

### nLFP detection and binning

For each condition, each electrode’s mean μ and standard deviation SD were computed over the retained samples. An nLFP was defined as a contiguous excursion of the LFP below μ − *z*·SD, timed at the most negative sample of the excursion, so that each excursion yields exactly one event; *z* = 3 unless stated otherwise, and *z* = 1.5–4.5 in steps of 0.5 for the threshold analysis. Event rasters at bin widths Δ*t* = 2, 4, 8, 16 and 32 ms were obtained by decimation, as in the original study: for Δ*t* = *n* × 2 ms the LFP was subsampled by keeping every *n*-th sample, starting with the first, and nLFPs were detected on the subsampled signal with the same μ and SD. Two alternatives were computed for comparison: detection of every local minimum below threshold, and detection at 2 ms followed by marking a wider bin as active if it contained at least one event (Supplementary Fig. S7). Phase averaging over the *n* possible start samples of the decimation was evaluated on one session and not applied otherwise (Results).

An avalanche is a maximal run of consecutive bins containing at least one nLFP, bounded by empty bins. Its size *s* is the total number of nLFPs in the run, summed over electrodes and bins, and its lifetime *T* is the number of bins.

### Exponent estimation

Size and lifetime exponents were estimated by discrete maximum likelihood [19] with the powerlaw package [20], with a fixed lower bound *x*_min_ = 1 for all bin widths and conditions. Fixing *x*_min_ rather than selecting it per fit by the Kolmogorov–Smirnov criterion is deliberate: a data-selected bound varies with Δ*t* and destabilises the α(Δ*t*) relation from which β is derived. The admissible exponent range was widened to 1–10 for lifetime and surrogate fits, because the package default (≤ 3) censors steep estimates at its boundary. The power law was compared with an exponential by the normalised log-likelihood ratio *R* (positive values favour the power law). Exponents are reported with the sign convention *P*(*s*) ∝ *s*^α^, *P*(*T*) ∝ *T*^τ^. Pooled fits combine the avalanches of all arrays and sessions of a group; per-file fits use one array–session. β was defined by α ∝ Δ*t*^β^ and estimated as the least-squares slope of log_10_|α| against log_10_ Δ*t* over 2–32 ms. Distributions are plotted with logarithmic binning.

### Branching ratio

Let *A*_t_ be the number of nLFPs in bin *t*. Three estimators were computed on each raster. (1) Within-avalanche: Σ*A*_t+1_/Σ*A*_t_ over pairs of consecutive non-empty bins within avalanches, excluding the terminating transition and avalanches of one bin. (2) Including terminations: Σ*A*_t+1_/Σ*A*_t_ over all bins with *A*_t_ > 0, so that the transition out of the last bin of an avalanche contributes zero descendants. (3) Beggs–Plenz [1]: for each avalanche, with *n*_a_ nLFPs in its first bin and *n*_d_ in the next (zero for one-bin avalanches), *d* = round(*n*_d_/*n*_a_); σ = Σ*n*_a_·*d*·(*N* − 1)/(*N* − *n*_a_) / Σ*n*_a_, where *N* is the number of electrodes and the factor corrects for refractoriness. Estimator (3) at Δ*t* = 2 ms is the one used in the source literature; all three are reported against Δ*t*, and estimator (2), which is the maximum-likelihood estimate of the mean offspring number of a branching process, is used for the statistical comparisons.

### Surrogates and controls

In the shuffled surrogate each electrode’s binned event sequence was independently permuted in time, preserving its event count while destroying the timing across electrodes; one surrogate per raster was generated and pooled across arrays. For the random-removal control, each array’s 1.5-SD raster at Δ*t* = 4 ms was thinned by deleting events uniformly at random down to the count obtained at each higher threshold, and avalanches were recomputed. For finite-size scaling, five random subsets of *N* = 12, 24, 36 and 48 electrodes were drawn from each array–session of monkey L with at least 48 retained electrodes; the cutoff was the 99th percentile of avalanche sizes, and the exponent *D* of cutoff ∝ *N*^D^ was fitted per array–session. For the cross-talk control, three random electrode sets per array, each as large as the full list removed, were compared with the list; all variants shared the same retained samples and per-electrode thresholds, so that they differed only in their electrodes. For duration matching, five windows per array, each as long as that array’s eyes-open recording, were drawn at random from the eyes-closed raster.

### Statistics

Bin widths within a file, eye states within an array and arrays within a session are not independent observations. Group comparisons therefore used linear mixed models (statsmodels MixedLM [21]) with fixed effects for the contrast of interest, log_2_ Δ*t* (centred at 4 ms) and their interaction, area as a covariate, and a variance component for arrays nested in sessions: parameter ∼ condition × log_2_ Δ*t* + area for eye state (monkey L) and parameter ∼ animal × log_2_ Δ*t* + area for the two animals (whole recordings). Models for β, which has one value per file, omit Δ*t*. Per-bin-width paired comparisons used Wilcoxon signed-rank tests with Benjamini–Hochberg correction [22]. Equivalence was assessed with two one-sided paired *t*-tests [10] and percentile-bootstrap confidence intervals of the mean paired difference (5,000 resamples of arrays). Confidence intervals of group means were obtained by bootstrapping arrays and those of pooled exponents by bootstrapping avalanches (2,000 resamples). Threshold slopes dα/d*z* were tested against zero, and against the random-removal slopes, with Wilcoxon signed-rank tests over array–sessions. Tests are two-sided with a significance level of 0.05 unless stated. The paired *t*-tests of the equivalence analysis were applied to 48 paired differences, for which the sampling distribution of the mean is approximately normal; the bootstrap confidence intervals, which make no normality assumption, led to the same conclusions.

Analyses were performed in Python 3.12.13 with NumPy 2.5.3 [23], SciPy 1.17.1 [24], pandas 2.3.3, powerlaw 2.0.0 [20], statsmodels 0.14.6 [21], matplotlib 3.10.8, seaborn 0.13.2 and statannotations 0.7.2.

Claude (Anthropic) was utilized solely for stylistic editing and language polishing of the manuscript. The authors reviewed and verified all content and assume full responsibility for the final text.

## Data availability

The recordings analysed are available from the G-Node Infrastructure, https://doi.org/10.12751/g-node.i20kyh [9]. We used, per session, the LFP files (NSP<n>_array<k>_LFP.mat), the eye-state epoch tables (epochs_<session>.csv), the cross-talk removal metadata tables, the SNR tables of the same-day checkerboard sessions (<SNR session>_full.csv) and the channel–area maps (channel_area_mapping_L.csv, channel_area_mapping_A.csv).

## Code availability

The repository contains the download scripts, the analysis library and the notebooks that generate every number and figure in this article.

## Acknowledgements

We thank X. Chen, P. R. Roelfsema and colleagues for making their recordings openly available. This work was supported by the MSU Institute for Artificial Intelligence.

## Author contributions

**K.S**.: Conceptualization, Methodology, Software, Investigation, Writing – original draft, Writing – review & editing.

**P.P**.: Software, Investigation, Data curation, Writing – original draft, Writing – review & editing.

**A.M**.: Formal analysis, Investigation, Writing – original draft (sections).

**M.L**.: Conceptualization, Project administration, Supervision, Writing – review & editing.

