## Supplementary Materials for "Neuronal avalanches in awake macaque V1 and V4"

Unless stated otherwise: one nLFP per below-threshold excursion, decimation binning, $z$ = 3 SD, arrays with ≥ 32 retained electrodes, monkey L with eyes-open and eyes-closed epochs analysed separately and monkey A as whole recordings. α, β and σ notation as in the main text.

#### Supplementary Figures

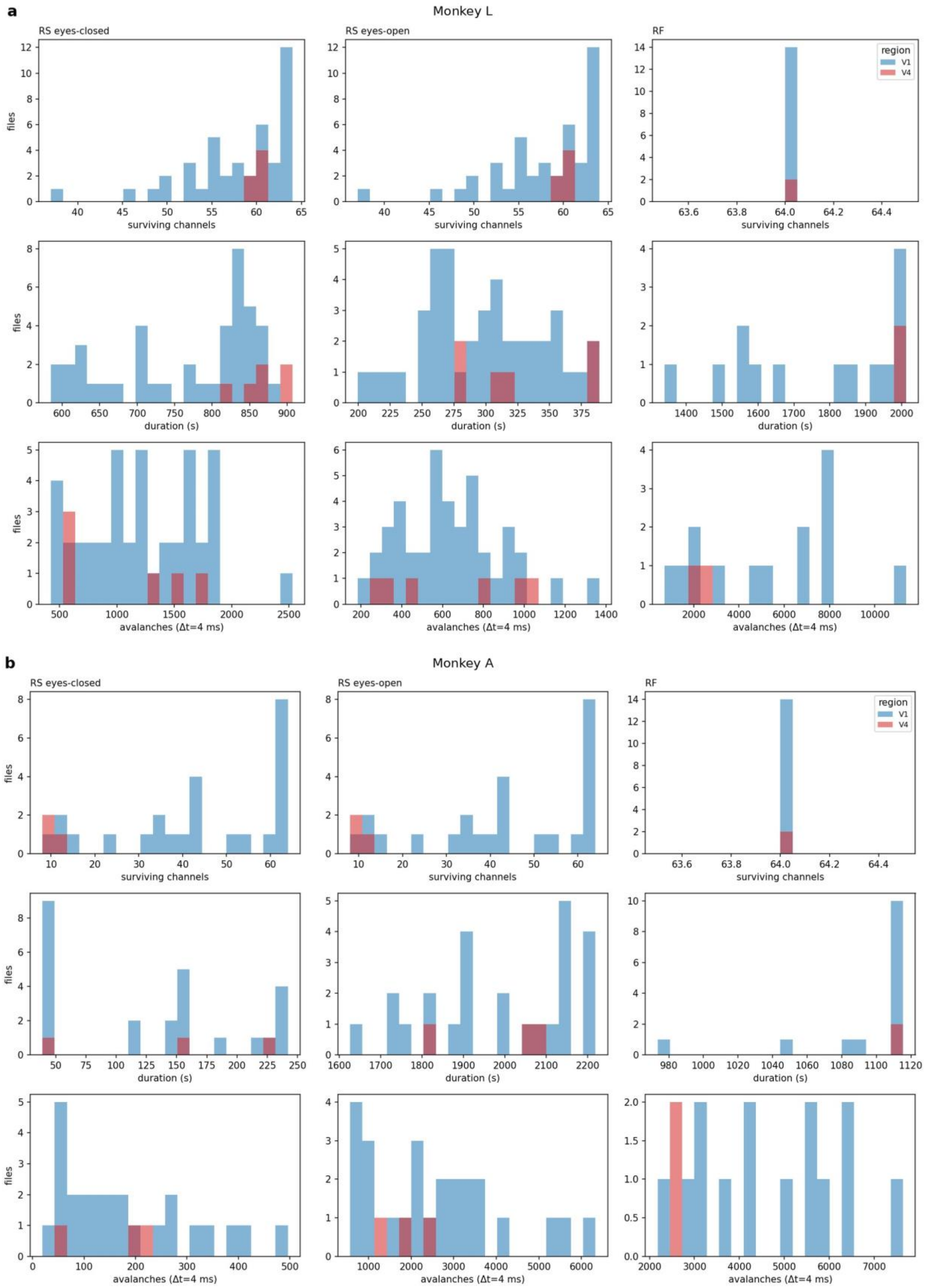


**Supplementary Figure S1. Data yield before reliability filtering.** Retained electrodes per array (top), condition duration after slow-wave exclusion (middle) and number of avalanches at $\Delta t$ = 4 ms (bottom), per array–session, for resting-state eyes closed, resting-state eyes open and the receptive-field mapping sessions (RF; not analysed in this study). V1 blue, V4 red. (**a**) Monkey L. (**b**) Monkey A; arrays with fewer than 32 retained electrodes, including both V4 arrays of this animal, were excluded from the analyses.

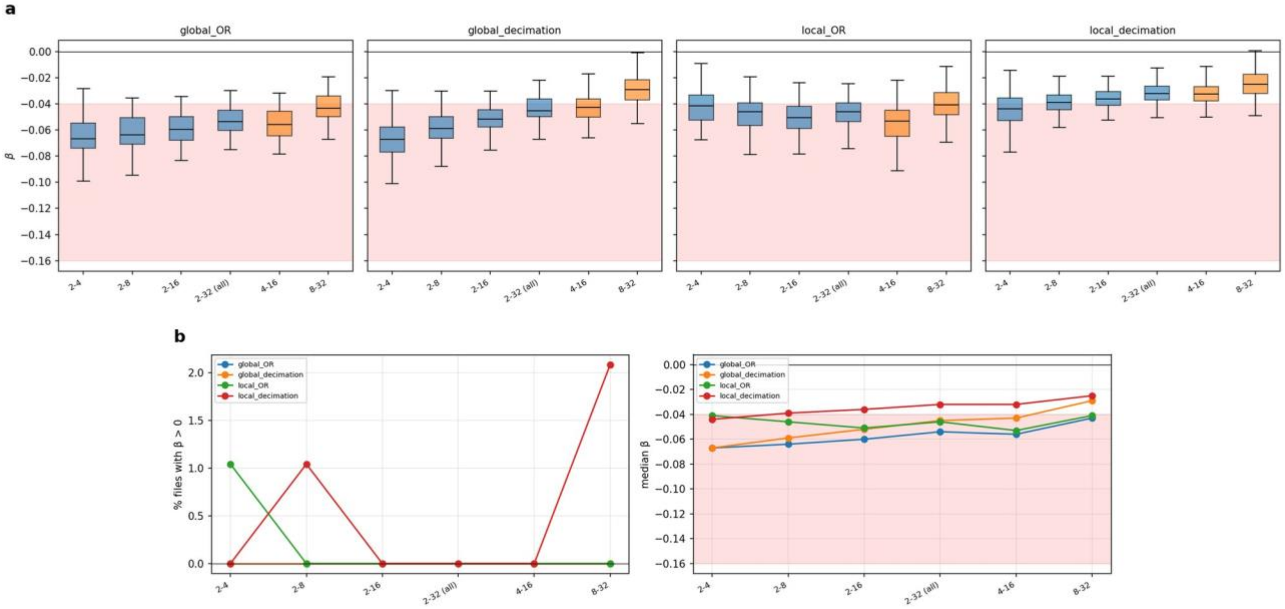


**Supplementary Figure S2. Dependence of β on the bin widths entering the fit** (monkey L, 96 array–session × eye-state files). (**a**) β per file for cumulative (blue: 2–4, 2–8, 2–16, 2–32 ms) and sliding (orange: 4–16, 8–32 ms) sets of bin widths, for the four combinations of event definition and binning (global = one event per excursion; local = every local minimum; decimation or OR binning). (**b**) Share of files with positive β and median β per set. Shaded band: range reported for M1/PMd (−0.16 to −0.04).

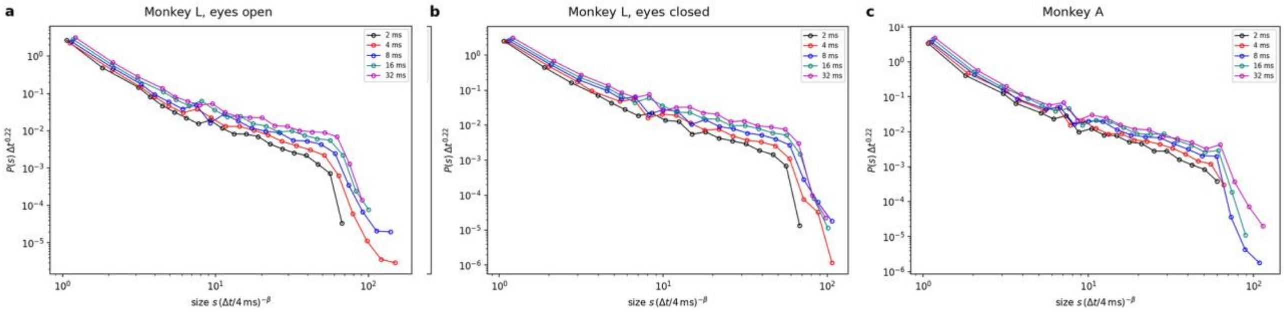


**Supplementary Figure S3. Scaling ansatz of the original study.** The distributions of Fig. 2a rescaled as $P(s)\cdot\Delta t^{0.22}$ against $s\cdot(\Delta t/4\text{ ms})^{-\beta}$, with the pooled β of each group (monkey L eyes open, −0.045; eyes closed, −0.046; monkey A, −0.031). The amplitude factor $\Delta t^{0.22}$ separates the bin widths rather than collapsing them; without it the distributions largely overlap (Fig. 2a).

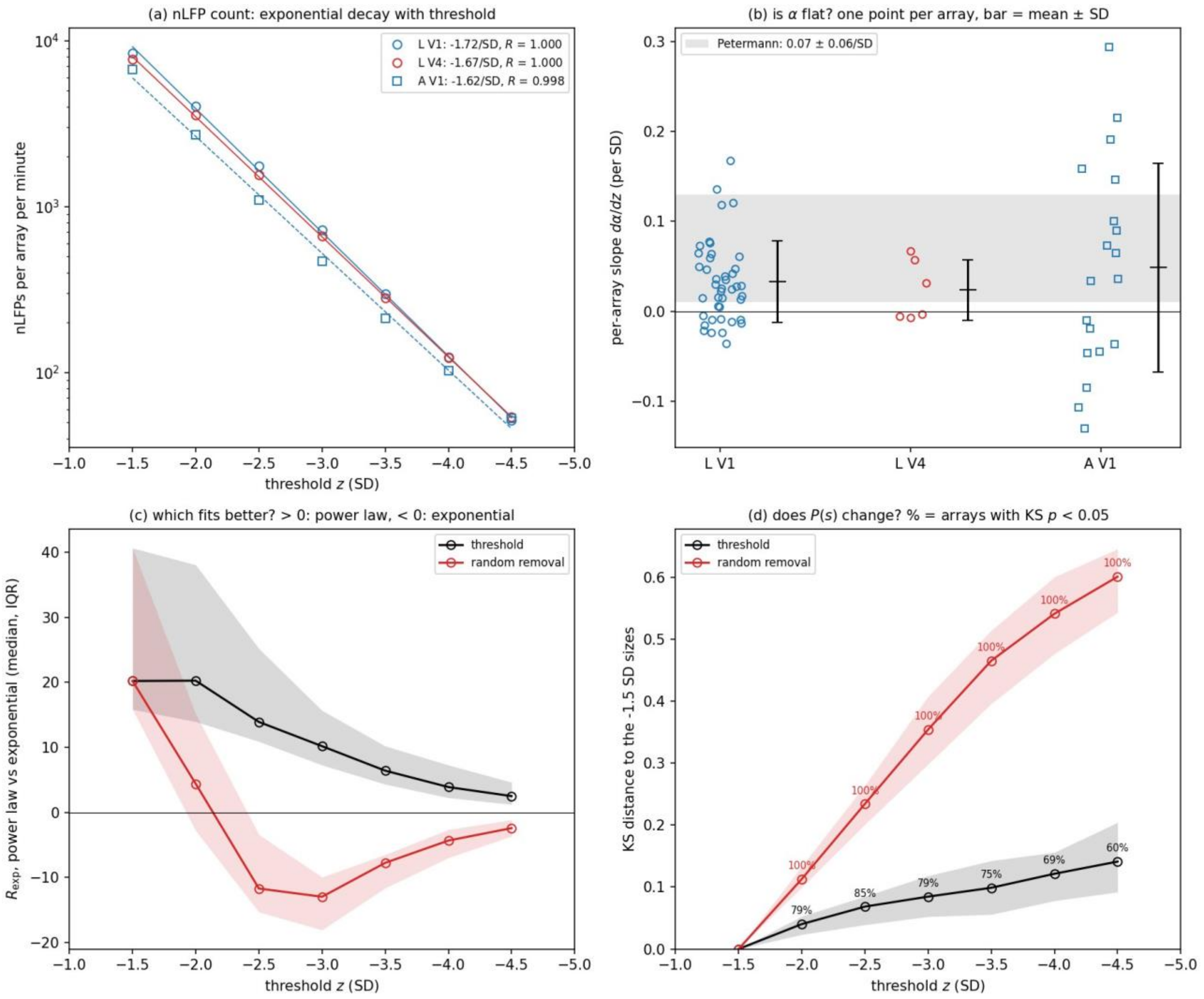


**Supplementary Figure S4. Threshold sweep summary** ($\Delta t$ = 4 ms, whole recordings; L V1 42, L V4 6, A V1 19 array–sessions). (**a**) nLFPs per array per minute on a logarithmic axis, with exponential fits. (**b**) Per-array slope dα/dz for threshold-based removal (mean ± SD), against the slope reported for motor cortex (shaded, 0.07 ± 0.06 per SD). (**c**) Median log-likelihood ratio of power law against exponential for threshold-based and random removal (IQR band). (**d**) Median Kolmogorov–Smirnov distance to the same array's 1.5-SD size distribution, with the share of array–sessions in which the difference is significant.

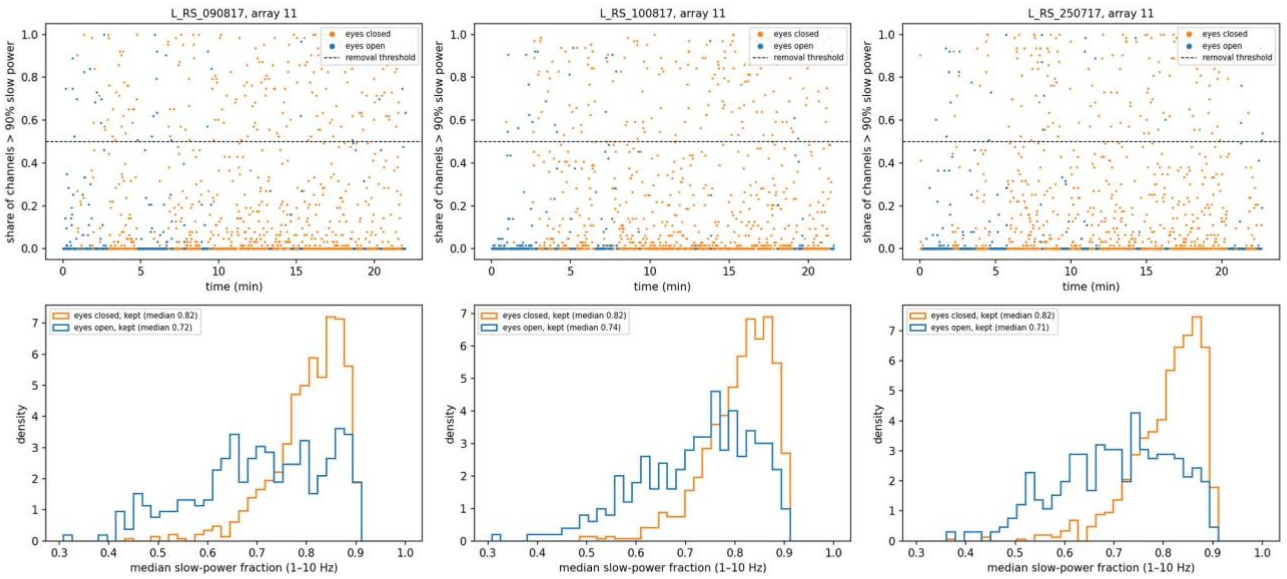


**Supplementary Figure S5. Slow-wave exclusion and residual slow activity in eyes-closed epochs** (array 11, each session of monkey L). Top: share of electrodes with more than 90% of their power in 1–10 Hz, per 2-s window, coloured by eye state; windows above 0.5 (line) are excluded. Bottom: distribution of the median slow-power fraction of retained eyes-open and eyes-closed windows.

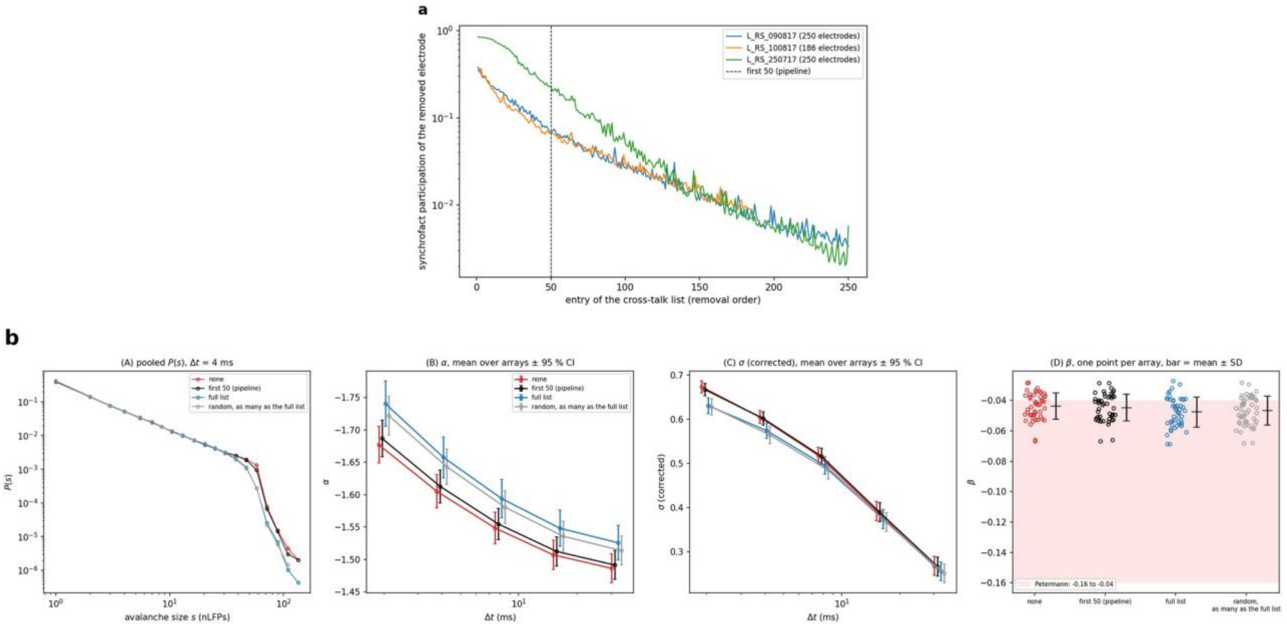


**Supplementary Figure S6. Cross-talk control** (monkey L). (**a**) Synchrofact participation along the ranked cross-talk removal list of each session; the pipeline removes the first 50 entries (line). (**b**) Pooled $P(s)$ at $\Delta t$ = 4 ms, α and σ against $\Delta t$ (mean ± 95% CI over array–sessions) and β per array–session, with no list, the first 50 entries (pipeline), the full list, and as many randomly chosen electrodes as the full list removed (three random sets per array).

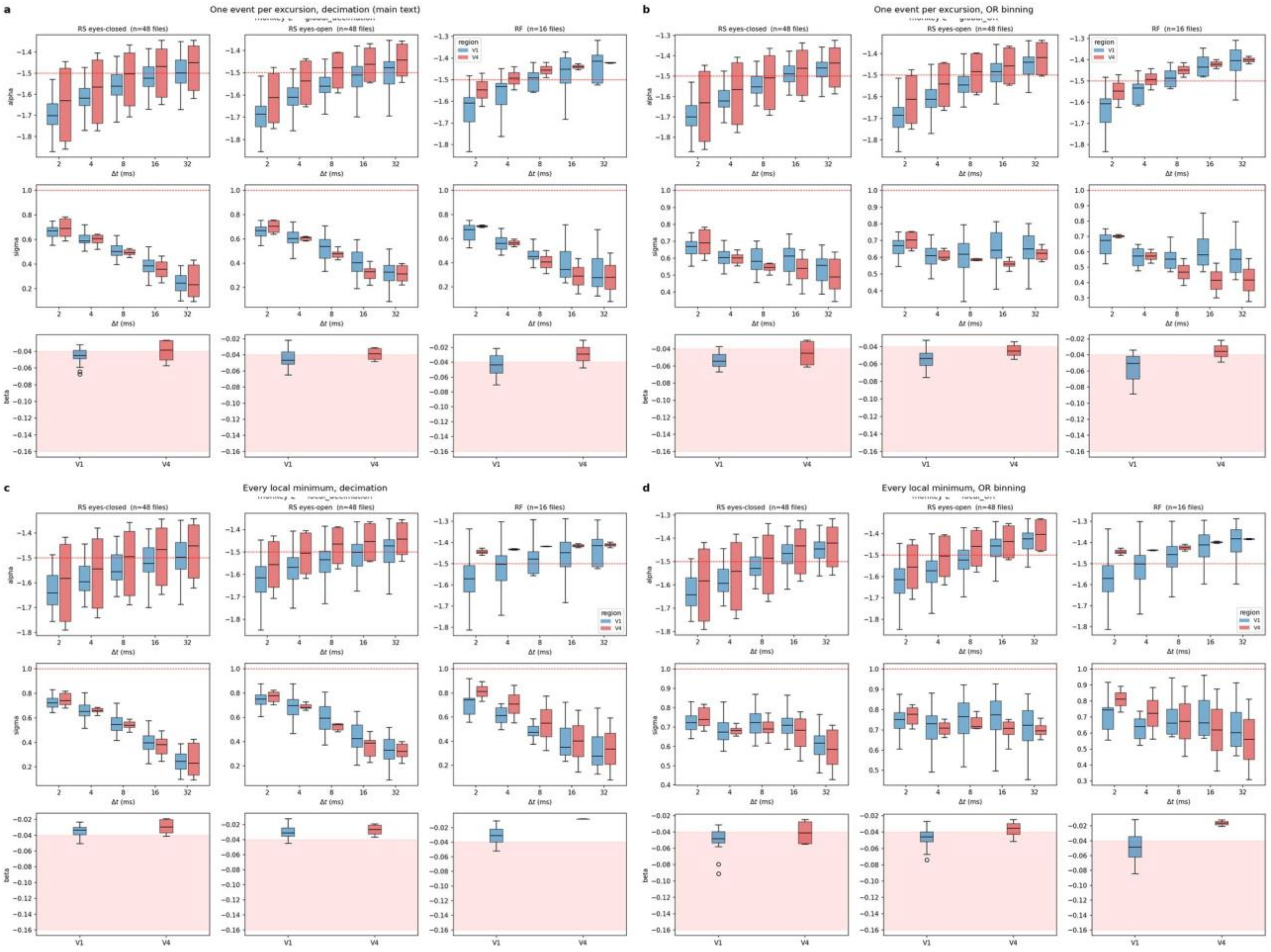


**Supplementary Figure S7. Event definition and binning** (monkey L). α and σ against $\Delta t$ and β, by condition (resting state eyes closed, eyes open; receptive-field mapping sessions, RF) and area (V1 blue, V4 red), for (**a**)

one event per below-threshold excursion with decimation (main text), (**b**) the same events with OR binning, (**c**) every local minimum below threshold with decimation and (**d**) with OR binning.

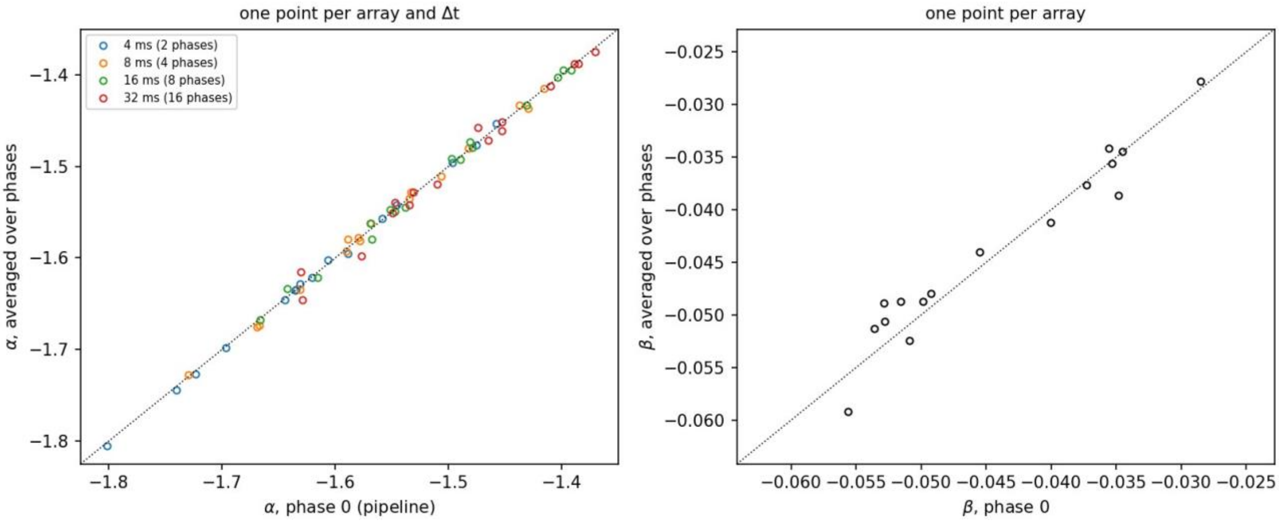


**Supplementary Figure S8. Phase averaging** (session L_RS_090817, 16 arrays). Left: α at decimation phase 0 against α of the distribution pooled over all $n = \Delta t/2$ ms phases, per array and bin width. Right: β from phase 0 against β from the phase-averaged exponents. Lines: identity.

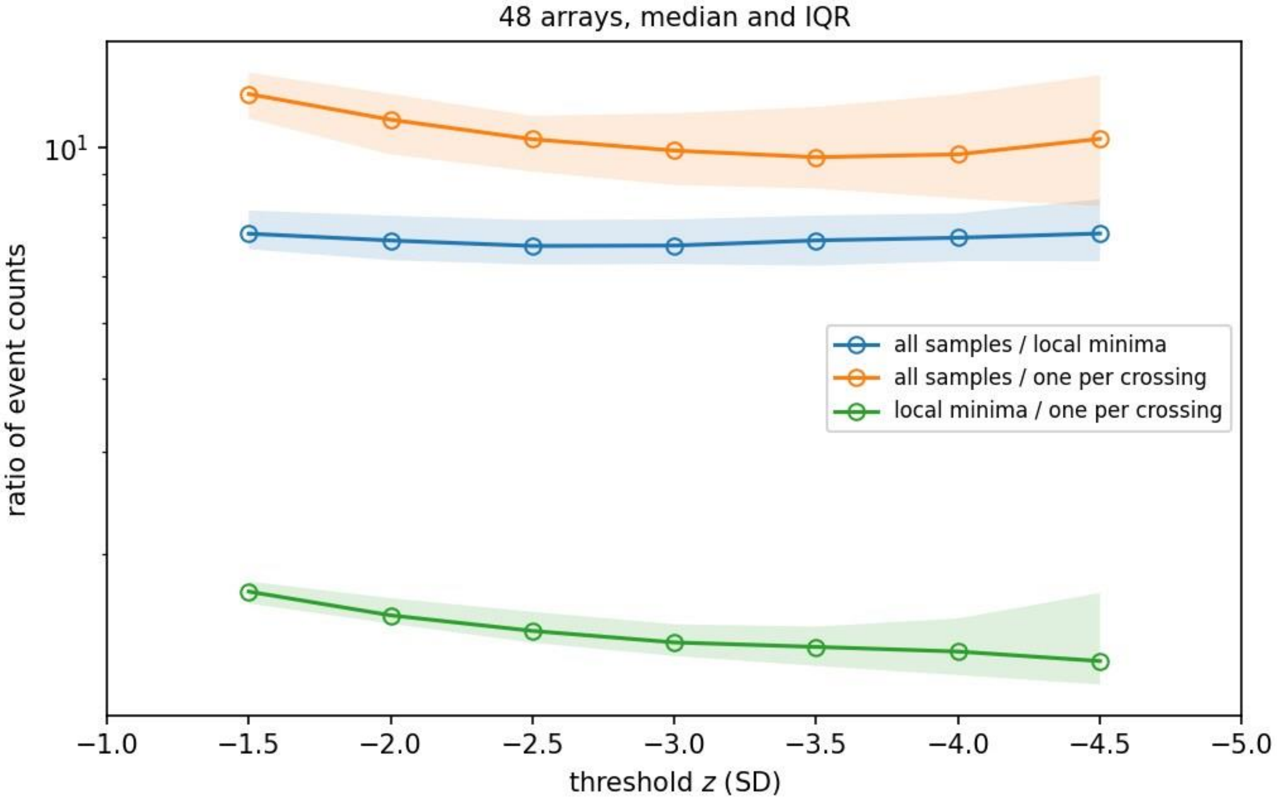


**Supplementary Figure S9. Event-count inflation.** Ratio of the number of below-threshold samples to the number of local minima below threshold (blue) and to the number of below-threshold excursions (orange), and ratio of local minima to excursions (green), against threshold; median and IQR over the 48 array–sessions of monkey L at 2-ms resolution.

### Supplementary Tables

**Table S1. Size exponents with bootstrap confidence intervals.** Mean over array–sessions with a 95% CI from resampling array–sessions (2,000 resamples), and the exponent of the pooled distribution with a 95% CI from resampling avalanches. The pooled CI describes how precisely one distribution is fitted; comparisons between groups should use the CI over arrays or the mixed models of Table S2.

| **Δ*t* (ms)** | **Arrays** | **Avalanches** | **α, mean over arrays** | **95% CI (arrays)** | **α, pooled** | **95% CI (avalanches)** |
|---|---|---|---|---|---|---|
| *L, eyes open* | | | | | | |
| 2 | 48 | 38,661 | −1.686 | −1.712 to −1.658 | −1.694 | −1.700 to −1.688 |
| 4 | 48 | 28,955 | −1.607 | −1.630 to −1.583 | −1.613 | −1.618 to −1.607 |
| 8 | 48 | 19,891 | −1.555 | −1.579 to −1.531 | −1.558 | −1.564 to −1.552 |
| 16 | 48 | 12,955 | −1.517 | −1.540 to −1.494 | −1.518 | −1.525 to −1.512 |
| 32 | 48 | 7,955 | −1.493 | −1.517 to −1.470 | −1.493 | −1.501 to −1.486 |
| *L, eyes closed* | | | | | | |
| 2 | 48 | 71,351 | −1.690 | −1.721 to −1.660 | −1.706 | −1.711 to −1.702 |
| 4 | 48 | 54,967 | −1.616 | −1.641 to −1.589 | −1.630 | −1.634 to −1.626 |
| 8 | 48 | 38,592 | −1.557 | −1.583 to −1.533 | −1.570 | −1.575 to −1.566 |
| 16 | 48 | 25,960 | −1.516 | −1.540 to −1.491 | −1.527 | −1.532 to −1.523 |
| 32 | 48 | 17,050 | −1.495 | −1.520 to −1.471 | −1.505 | −1.510 to −1.500 |
| *A, whole recordings* | | | | | | |
| 2 | 19 | 68,302 | −1.790 | −1.876 to −1.712 | −1.822 | −1.828 to −1.816 |
| 4 | 19 | 56,257 | −1.725 | −1.804 to −1.657 | −1.755 | −1.761 to −1.749 |
| 8 | 19 | 42,531 | −1.676 | −1.752 to −1.608 | −1.704 | −1.710 to −1.698 |
| 16 | 19 | 30,485 | −1.648 | −1.721 to −1.581 | −1.675 | −1.682 to −1.669 |
| 32 | 19 | 20,389 | −1.649 | −1.726 to −1.574 | −1.673 | −1.681 to −1.664 |

**Table S2. Linear mixed models.** Fixed-effect estimates with 95% CIs from models with a variance component for arrays nested in sessions (Methods); effects are evaluated at Δ*t* = 4 ms ($\log_2$ Δ*t* centred there) or per doubling of Δ*t*. The last column gives the *P* value of the same formula fitted by ordinary least squares with every row treated as independent. For β, which has one value per file, the models omit Δ*t*; in the animal model each array then contributes one row per session and the array component is replaced by a session intercept.

| | **Effect** | **Estimate** | **95% CI** | ***P* (mixed)** | ***P* (rows independent)** |
|---|---|---|---|---|---|
| *Eye state (monkey L, 48 array–sessions × 2 eye states × 5 Δt)* | | | | | |
| α | Eyes closed − open | −0.0049 | −0.0129 to 0.0031 | 0.23 | 0.62 |
| α | V4 − V1 | 0.0528 | −0.0180 to 0.1235 | 0.14 | $1.9 \times 10^{-5}$ |
| α | Per doubling of Δ*t* | 0.0476 | 0.0443 to 0.0509 | $1 \times 10^{-178}$ | $3 \times 10^{-28}$ |
| α | Closed − open, per doubling of Δ*t* | 0.0015 | −0.0031 to 0.0061 | 0.53 | 0.79 |
| σ | Eyes closed − open | −0.0050 | −0.0184 to 0.0085 | 0.47 | 0.58 |
| σ | V4 − V1 | −0.0108 | −0.0575 to 0.0358 | 0.65 | 0.32 |
| σ | Per doubling of Δ*t* | −0.0910 | −0.0965 to −0.0855 | $1 \times 10^{-230}$ | $3 \times 10^{-89}$ |
| σ | Closed − open, per doubling of Δ*t* | −0.0155 | −0.0233 to −0.0078 | $9 \times 10^{-5}$ | 0.0025 |
| β | Eyes closed − open | −0.0012 | −0.0039 to 0.0016 | 0.40 | 0.56 |
| β | V4 − V1 | 0.0052 | −0.0024 to 0.0127 | 0.18 | 0.098 |
| *Animals (whole recordings; monkey L 48, monkey A 19 array–sessions × 5 Δt)* | | | | | |
| α | Monkey A − L | −0.107 | −0.170 to −0.044 | $8.8 \times 10^{-4}$ | $2.9 \times 10^{-9}$ |
| α | V4 − V1 | 0.047 | −0.053 to 0.146 | 0.36 | 0.041 |
| α | Per doubling of Δ*t* | 0.0491 | 0.0469 to 0.0514 | $< 10^{-300}$ | $3 \times 10^{-18}$ |
| α | A − L, per doubling of Δ*t* | −0.0133 | −0.0175 to −0.0091 | $6.0 \times 10^{-10}$ | 0.18 |
| σ | Monkey A − L | −0.0438 | −0.0754 to −0.0123 | 0.0064 | $7.0 \times 10^{-5}$ |

| | Effect | Estimate | 95% CI | *P* (mixed) | *P* (rows independent) |
|---|---|---|---|---|---|
| σ | V4 − V1 | −0.0102 | −0.0580 to 0.0376 | 0.67 | 0.47 |
| σ | Per doubling of Δ*t* | −0.1014 | −0.1061 to −0.0967 | $< 10^{-300}$ | $9 \times 10^{-99}$ |
| σ | A − L, per doubling of Δ*t* | −0.0042 | −0.0130 to 0.0046 | 0.35 | 0.50 |
| β | Monkey A − L | 0.0149 | 0.0099 to 0.0198 | $4.0 \times 10^{-9}$ | $6.6 \times 10^{-8}$ |
| β | V4 − V1 | 0.0042 | −0.0033 to 0.0117 | 0.27 | 0.28 |
| *Variance components* | | | | | |
| α | Eye state model | array SD 0.082 | residual SD 0.037 | array share 83% | |
| σ | Eye state model | array SD 0.051 | residual SD 0.062 | array share 41% | |
| β | Eye state model | array SD 0.0073 | residual SD 0.0068 | array share 53% | |
| α | Animals model | array SD 0.116 | residual SD 0.025 | array share 95% | |
| σ | Animals model | array SD 0.051 | residual SD 0.052 | array share 48% | |

**Table S3. Threshold sweep and random-removal control** (Δ*t* = 4 ms, whole recordings). Per threshold *z*: nLFPs per array per minute; α as the mean over array–sessions with a 95% bootstrap CI over array–sessions; mean avalanche size ⟨*s*⟩; median normalised log-likelihood ratio *R* of power law against exponential (positive favours the power law); median Kolmogorov–Smirnov distance *D* to the same array's 1.5-SD size distribution. Random removal deletes 1.5-SD events at random down to the event count of each threshold.

| | | Threshold-based removal | | | | Random removal | | | |
|---|---|---|---|---|---|---|---|---|---|
| *z* (SD) | nLFP $min^{-1}$ | α [95% CI] | ⟨*s*⟩ | *R* | *D* | α [95% CI] | ⟨*s*⟩ | *R* | *D* |
| *Monkey L, V1 (42 array–sessions)* | | | | | | | | | |
| −1.5 | 8,404 | −1.54 [−1.56, −1.53] | 9.5 | 16.7 | 0.00 | −1.54 [−1.56, −1.53] | 9.5 | 16.7 | 0.00 |
| −2.0 | 4,032 | −1.57 [−1.59, −1.56] | 8.6 | 16.8 | 0.04 | −1.65 [−1.67, −1.64] | 5.4 | 1.4 | 0.11 |
| −2.5 | 1,761 | −1.60 [−1.62, −1.58] | 8.0 | 12.3 | 0.06 | −1.85 [−1.88, −1.82] | 3.1 | −11.6 | 0.24 |
| −3.0 | 726 | −1.62 [−1.64, −1.60] | 7.6 | 8.7 | 0.07 | −2.21 [−2.27, −2.15] | 1.9 | −12.2 | 0.36 |
| −3.5 | 298 | −1.62 [−1.65, −1.60] | 7.5 | 5.3 | 0.08 | −2.80 [−2.91, −2.70] | 1.4 | −7.6 | 0.47 |
| −4.0 | 123 | −1.63 [−1.67, −1.59] | 7.6 | 3.2 | 0.10 | −3.67 [−3.85, −3.48] | 1.2 | −3.9 | 0.56 |
| −4.5 | 51 | −1.65 [−1.70, −1.61] | 7.2 | 1.8 | 0.11 | −4.71 [−5.01, −4.44] | 1.1 | −1.9 | 0.62 |
| *Monkey L, V4 (6 array–sessions)* | | | | | | | | | |
| −1.5 | 7,747 | −1.52 [−1.58, −1.44] | 11.4 | 11.4 | 0.00 | −1.52 [−1.58, −1.45] | 11.4 | 11.4 | 0.00 |
| −2.0 | 3,548 | −1.54 [−1.62, −1.46] | 10.7 | 12.7 | 0.03 | −1.64 [−1.73, −1.54] | 6.1 | −2.7 | 0.16 |
| −2.5 | 1,549 | −1.55 [−1.64, −1.46] | 10.6 | 8.5 | 0.03 | −1.83 [−1.96, −1.70] | 3.4 | −12.9 | 0.28 |
| −3.0 | 666 | −1.57 [−1.68, −1.46] | 10.5 | 6.6 | 0.05 | −2.15 [−2.36, −1.95] | 2.1 | −16.1 | 0.41 |
| −3.5 | 280 | −1.58 [−1.70, −1.47] | 10.4 | 5.2 | 0.08 | −2.68 [−2.98, −2.38] | 1.5 | −9.5 | 0.50 |
| −4.0 | 123 | −1.58 [−1.70, −1.46] | 10.7 | 2.6 | 0.06 | −3.42 [−3.85, −3.01] | 1.2 | −5.1 | 0.58 |
| −4.5 | 54 | −1.59 [−1.75, −1.45] | 11.3 | 2.0 | 0.13 | −4.30 [−4.88, −3.79] | 1.1 | −3.1 | 0.63 |
| *Monkey A, V1 (19 array–sessions)* | | | | | | | | | |
| −1.5 | 6,730 | −1.63 [−1.66, −1.60] | 7.4 | 49.5 | 0.00 | −1.63 [−1.66, −1.60] | 7.4 | 49.5 | 0.00 |
| −2.0 | 2,701 | −1.71 [−1.77, −1.66] | 6.3 | 39.9 | 0.06 | −1.77 [−1.83, −1.73] | 4.0 | 9.1 | 0.11 |
| −2.5 | 1,095 | −1.74 [−1.81, −1.68] | 6.2 | 30.4 | 0.10 | −2.02 [−2.11, −1.94] | 2.4 | −12.3 | 0.22 |
| −3.0 | 467 | −1.72 [−1.80, −1.66] | 6.8 | 19.3 | 0.12 | −2.42 [−2.57, −2.28] | 1.7 | −15.1 | 0.33 |
| −3.5 | 212 | −1.72 [−1.82, −1.64] | 7.2 | 10.3 | 0.14 | −3.00 [−3.25, −2.75] | 1.4 | −9.9 | 0.42 |

| | | | | | | | | | |
|---|---|---|---|---|---|---|---|---|---|
| −4.0 | 102 | −1.76 [−1.87, −1.66] | 7.0 | 7.8 | 0.15 | −3.74 [−4.15, −3.34] | 1.2 | −5.7 | 0.49 |
| −4.5 | 53 | −1.83 [−1.99, −1.69] | 6.7 | 5.5 | 0.18 | −4.54 [−5.14, −3.96] | 1.1 | −3.6 | 0.54 |

**Table S4. Branching ratio.** Mean ± SD over array–sessions for the estimator including terminations, the Beggs–Plenz estimator and the within-avalanche estimator (Methods), and the estimator including terminations on per-electrode shuffled rasters (monkey L; for monkey A see Fig. 5a).

| Δ*t* (ms) | Including terminations | Beggs–Plenz | Within-avalanche | Shuffled (incl. terminations) |
|---|---|---|---|---|
| *Monkey L, eyes open (48 array–sessions)* | | | | |
| 2 | 0.67 ± 0.05 | 0.95 ± 0.29 | 0.99 ± 0.03 | 0.030 |
| 4 | 0.61 ± 0.07 | 0.82 ± 0.13 | 1.00 ± 0.05 | 0.057 |
| 8 | 0.52 ± 0.09 | 0.62 ± 0.16 | 0.99 ± 0.10 | 0.095 |
| 16 | 0.39 ± 0.11 | 0.42 ± 0.17 | 0.99 ± 0.14 | 0.137 |
| 32 | 0.32 ± 0.10 | 0.34 ± 0.15 | 1.02 ± 0.27 | 0.180 |
| *Monkey L, eyes closed (48 array–sessions)* | | | | |
| 2 | 0.67 ± 0.05 | 0.96 ± 0.28 | 0.99 ± 0.02 | 0.021 |
| 4 | 0.60 ± 0.06 | 0.82 ± 0.17 | 1.00 ± 0.03 | 0.040 |
| 8 | 0.51 ± 0.06 | 0.62 ± 0.11 | 1.00 ± 0.06 | 0.068 |
| 16 | 0.38 ± 0.08 | 0.45 ± 0.12 | 1.01 ± 0.12 | 0.104 |
| 32 | 0.24 ± 0.08 | 0.26 ± 0.11 | 1.00 ± 0.23 | 0.144 |
| *Monkey A, whole recordings (19 array–sessions)* | | | | |
| 2 | 0.64 ± 0.09 | 1.04 ± 0.38 | 0.99 ± 0.01 | — |
| 4 | 0.55 ± 0.07 | 0.78 ± 0.21 | 0.98 ± 0.03 | — |
| 8 | 0.45 ± 0.08 | 0.58 ± 0.15 | 0.97 ± 0.03 | — |
| 16 | 0.33 ± 0.09 | 0.40 ± 0.15 | 0.98 ± 0.08 | — |
| 32 | 0.22 ± 0.09 | 0.20 ± 0.08 | 0.90 ± 0.11 | — |

**Table S5. Eye state in monkey L.** Top: mean paired difference (eyes closed − eyes open) per bin width, Wilcoxon signed-rank *P* and Benjamini–Hochberg-corrected *P*. Bottom: mean paired difference with 95% bootstrap CI over array–sessions (5,000 resamples), *P* of the two one-sided tests (TOST) against the stated equivalence bound, and the Wilcoxon *P* of the difference. Bounds: one third of the range of the parameter across arrays (α 0.117, σ 0.068, β 0.009) and, for α, a stricter 0.07. Duration-matched: eyes-closed parameters averaged over five random windows per array of the eyes-open duration.

| | Comparison | Mean difference | 95% CI | *P* (Wilcoxon) / *P* (TOST) | $P_{BH}$ / *P* (Wilcoxon) |
|---|---|---|---|---|---|
| *Paired Wilcoxon signed-rank tests per bin width (48 array–sessions)* | | | | | |
| α | Δ*t* = 2 ms | −0.0046 | | 0.27 | 0.68 |
| α | Δ*t* = 4 ms | −0.0092 | | 0.17 | 0.68 |
| α | Δ*t* = 8 ms | −0.0019 | | 0.87 | 0.87 |
| α | Δ*t* = 16 ms | 0.0009 | | 0.69 | 0.87 |
| α | Δ*t* = 32 ms | −0.0022 | | 0.60 | 0.87 |
| σ | Δ*t* = 2 ms | −0.0001 | | 0.99 | 0.99 |
| σ | Δ*t* = 4 ms | −0.0063 | | 0.34 | 0.43 |
| σ | Δ*t* = 8 ms | −0.0088 | | 0.35 | 0.43 |
| σ | Δ*t* = 16 ms | −0.0129 | | 0.29 | 0.43 |
| σ | Δ*t* = 32 ms | −0.0745 | | $5.4 \times 10^{-8}$ | $2.7 \times 10^{-7}$ |
| *Equivalence tests (two one-sided paired* t*-tests; Δt = 4 ms for α and σ)* | | | | | |

| | Comparison | Mean difference | 95% CI | *P* (Wilcoxon) / *P* (TOST) | $P_{BH}$ / *P* (Wilcoxon) |
|---|---|---|---|---|---|
| α | Original, bound ±0.070 | −0.0092 | −0.0243 to 0.0050 | $7.0 \times 10^{-11}$ | 0.17 |
| α | Original, bound ±0.117 | −0.0092 | −0.0243 to 0.0050 | $3.0 \times 10^{-19}$ | 0.17 |
| α | Duration-matched, bound ±0.070 | −0.0256 | −0.0410 to −0.0102 | $6.5 \times 10^{-7}$ | 0.0042 |
| α | Duration-matched, bound ±0.117 | −0.0256 | −0.0410 to −0.0102 | $1.9 \times 10^{-15}$ | 0.0042 |
| σ | Original, bound ±0.068 | −0.0063 | −0.0177 to 0.0048 | $3.6 \times 10^{-14}$ | 0.34 |
| σ | Duration-matched, bound ±0.068 | −0.0161 | −0.0283 to −0.0041 | $7.5 \times 10^{-11}$ | 0.012 |
| β | Original, bound ±0.009 | −0.0012 | −0.0039 to 0.0015 | $1.6 \times 10^{-7}$ | 0.52 |
| β | Duration-matched, bound ±0.009 | −0.0022 | −0.0049 to 0.0005 | $1.8 \times 10^{-6}$ | 0.27 |
